# Neuromedin U signaling regulates memory retrieval of learned salt avoidance in a *C. elegans* gustatory circuit

**DOI:** 10.1101/683888

**Authors:** Jan Watteyne, Petrus Van der Auwera, Katleen Peymen, Charline Borghgraef, Elke Vandewyer, Iene Rutten, Jeroen Lammertyn, Rob Jelier, Liliane Schoofs, Isabel Beets

## Abstract

Learning and memory are regulated by neuromodulatory pathways, but the contribution and temporal requirement of most neuromodulators in a learning circuit are unknown. Here we identify the evolutionarily conserved neuromedin U (NMU) neuropeptide family as a regulator of memory retrieval in *C. elegans* gustatory aversive learning. The NMU homolog CAPA-1 and its receptor NMUR-1 are required for the expression of learned salt avoidance. Aversive learning depends on the release of CAPA-1 neuropeptides from sensory ASG neurons that respond to salt stimuli in an experience-dependent manner. Optogenetic silencing of CAPA-1 neurons blocks the immediate retrieval, but not the acquisition, of learned salt avoidance. CAPA-1 subsequently signals through NMUR-1 in AFD sensory neurons to modulate two navigational strategies for salt chemotaxis. Aversive conditioning thus recruits NMU signaling to eventually modulate locomotor programs for expressing learned avoidance behavior. Because NMU signaling is conserved across bilaterian animals, our findings incite further research into its function in other memory and decision-making circuits.

**Graphical Abstract:** 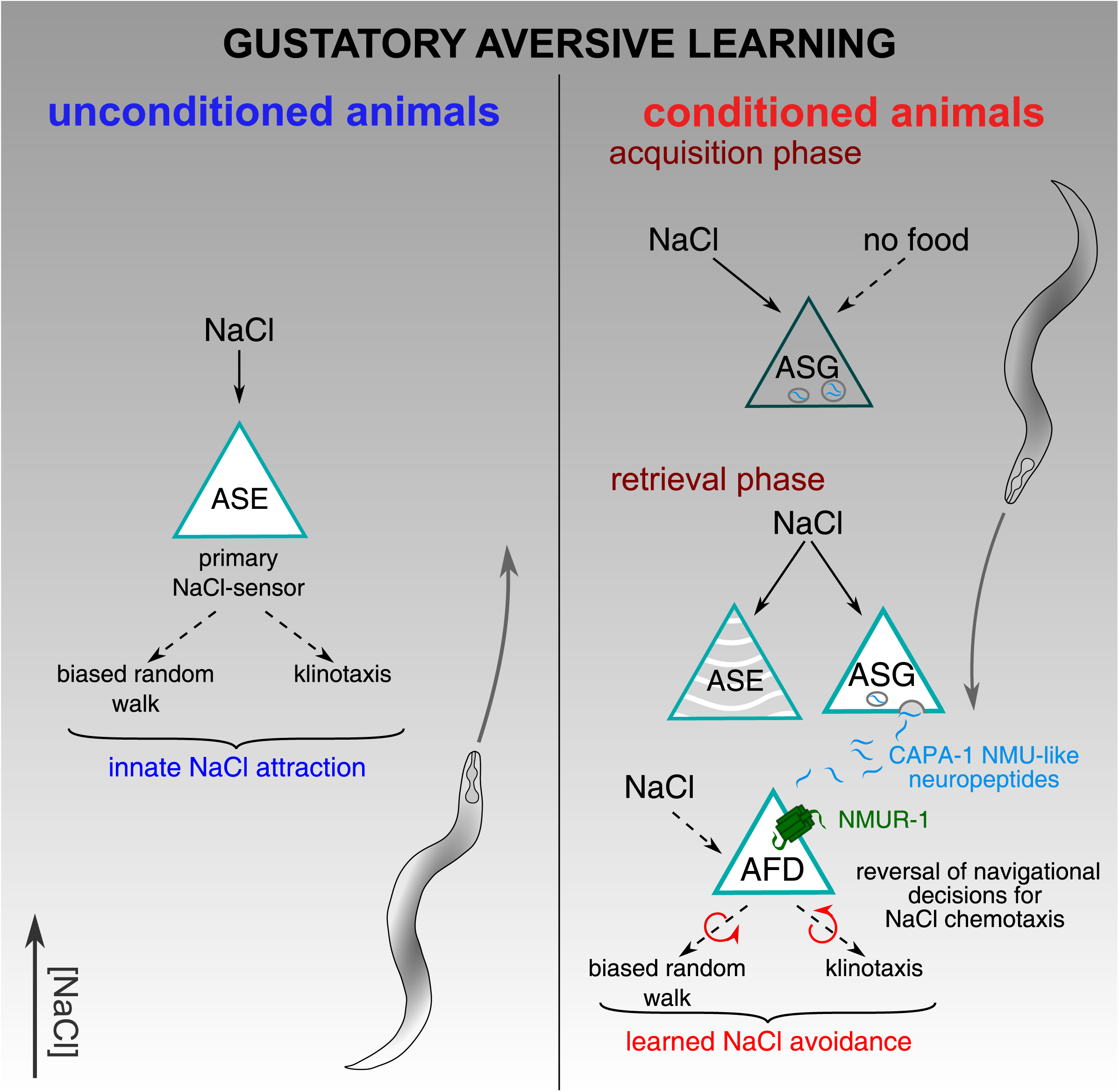

## Introduction

Associative learning and memories are crucial for animals to anticipate events and adapt behavioral choices, increasing their chance of survival in a dynamic environment. The experience-dependent association of two stimuli, known as Pavlovian conditioning, creates memory traces that can profoundly influence future behavioral responses^1^. Early studies in invertebrate model organisms, such as *Aplysia*, have been key to provide insights in neural mechanisms underlying learning and memory^2,3^. Since then it has been suggested in diverse model systems that molecular components of associative learning seem to be evolutionarily conserved from worms to humans^4^. Since memories, in particular of negative experiences, have a large impact on decision-making as well as cognitive and emotional states, mechanisms underlying these functions may share common elements.

Modifications in neural signaling mediate and maintain experience-driven changes in brain circuits for memory^5^. The hypothesis that neuropeptides are central modulators of such learning circuits is currently emerging, and neuropeptide receptors have gained significant interest as potential key targets for treatment of cognitive disorders^6–10^. Neuropeptides make up an evolutionary ancient and diverse class of neural messengers that serve a broad range of functions^11–15^. They modulate neural signaling mainly by binding G protein-coupled receptors (GPCRs)^16,17^ and are widely distributed in brain areas responsible for learning and memory^18–21^.

A role in experience-dependent plasticity has been established for some neuropeptides, but many questions remain concerning the actions of neuromodulators in learning circuits. Behaviorally, it is unclear whether they regulate specific aspects of a learned behavior or serve as general modulators that coordinately control distinct conditioned responses to a stimulus. Also, the temporal requirement of peptidergic neurons in learning circuits is largely unexplored, in part because most experimental studies examined the effects of peptide injections into the brain, which is limited in temporal and spatial resolution for studying neuropeptidergic modulation^18,20,21^.

With a nervous system of 302 neurons, *Caenorhabditis elegans* provides an excellent model system for dissecting learning and neuropeptidergic circuits^22–26^. The physical connectivity of neurons has been mapped and the nematode shows different kinds of plasticity in gustatory, olfactory, thermo- and mechanosensory behaviors. These include associative learning and memory, which require molecular mechanisms similar to those in vertebrates^11,27,28^. We previously found that gustatory associative learning in *C. elegans* is regulated by nematocin, a neuropeptide of the vasopressin/oxytocin family that promotes salt aversion when worms experienced salt in the absence of food^10^. The *C. elegans* genome counts at least 153 neuropeptide-encoding genes and 150 peptide GPCRs^29–31^, many of which are evolutionarily conserved between worms and humans^32–34^. Learning is likely regulated by multiple neuromodulators, yet our understanding of how individual modulators contribute to a learned behavior in a defined circuit is limited.

To uncover neuropeptidergic pathways that regulate learning circuits, we tested *C. elegans* mutants of neuropeptide receptors for their performance in gustatory associative learning. We focused on evolutionarily conserved neuropeptide systems that are expressed in brain regions for learning and memory. Such a neuropeptide system is the neuromedin U (NMU) pathway that is present in most bilaterian animals^32,35^. We hypothesized that NMU-like signaling regulates learning because NMU peptides and their receptors are widely expressed in memory centers, such as in the vertebrate hippocampus and the insect mushroom bodies^36–38^. In mammals, intranasal administration or injection of NMU peptides into the brain affects memory and reward-related behaviors, further suggesting a role for NMU in experience-dependent plasticity^39–43^, although mechanisms underlying these effects remain unknown.

Here, we demonstrate that NMU neurons are specifically required for the retrieval of learned salt aversion in *C. elegans*. In addition, we show that NMU neuropeptides released from a pair of sensory neurons, i.e. the ASG cells that show experience-dependent responses to salt, signal via their receptor NMUR-1 on the AFD sensory neurons to coordinately regulate distinct navigational decisions for adapting salt chemotaxis behavior. While it is widely assumed in many fields, including neuropsychology and neuroeconomics, that decision-making involves learning and memory, the underlying molecular underpinnings and neural mechanisms are still not well understood. Our findings indicate a role for the NMU neuropeptide family as a modulator of the expression of learned aversion, and lay a foundation for further research examining the functions of the conserved NMU system in other learning circuits underlying decision-making tasks.

## Results

### A *C. elegans* neuromedin receptor ortholog, NMUR-1, regulates gustatory aversive learning

*C. elegans* has three predicted NMU receptor orthologs, named NMUR-1 to NMUR-3^32,34^ (Figure 1a). We hypothesized that this receptor family is involved in experience-dependent plasticity, because NMU receptors are widely expressed in memory centers of the brain^36–38,44^. To test this hypothesis, we analyzed the performance of *nmur* loss-of-function mutants (Figure 1b) in an established paradigm for gustatory associative learning^10,45^. Similar to classical aversive conditioning, we trained worms by pairing a conditioned salt stimulus with the absence of food during a 15-minute training period (Figure 1c). We then tested salt chemotaxis behavior by putting worms in the center of a quadrant plate and calculated a chemotaxis index based on the number of animals that migrated towards or away from NaCl-containing quadrants (Figure 1c). Mock-conditioned worms, which have not been pre-exposed to salt, were strongly attracted to NaCl and wild-type animals learned to avoid salt when they previously experienced it in the absence of food (Figure 1d). Consistent with our hypothesis, *nmur-1* mutants were defective in learning salt aversion and were still attracted to salt after aversive NaCl-conditioning (Figure 1d). Mutants for the NMU receptor orthologs *nmur-2* and *nmur-3* had no plasticity defects (Figure 1d), suggesting a role specific for NMUR-1 in gustatory aversive learning.

**Figure 1.**
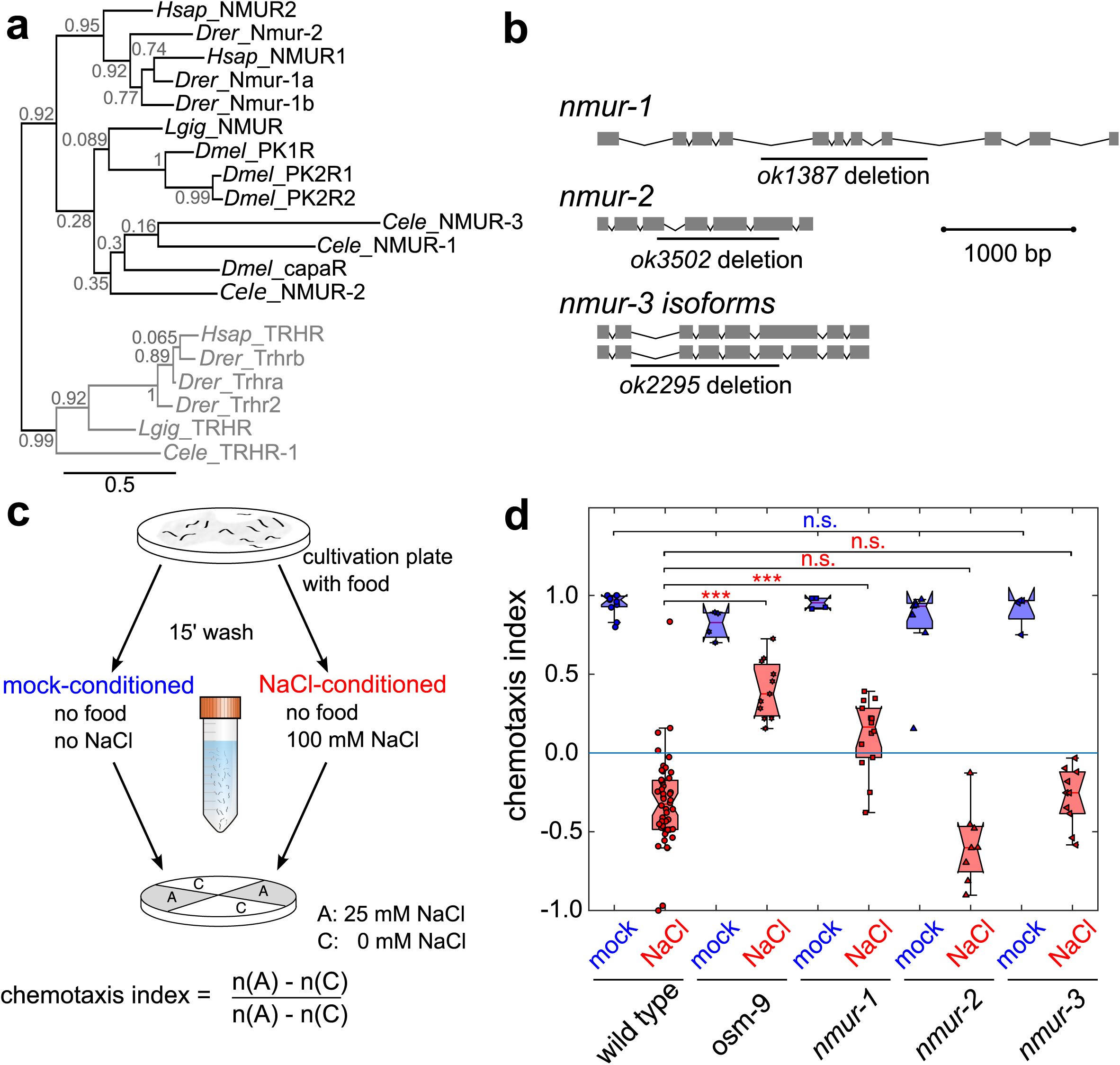
The *C. elegans* Neuromedin U receptor ortholog NMUR-1 regulates gustatory associative learning (a) Maximum likelihood tree of the phylogenetic relationship between neuromedin U receptor (NMUR) orthologs of *C. elegans* (*Cele*), *D. melanogaster* (*Dmel*), *L. gigantean* (*Lgig*), *D. rerio* (*D. rerio*) and human (*Hsap*). Thyrotropin-releasing hormone receptors (TRHRs) are used as outgroup to root the tree, because this family of receptors was previously shown to be most closely related to bilaterian NMURs^32^. Branch node labels correspond to likelihood ratio test values. **(b)** Gene organization of the three predicted *nmur* orthologs in *C. elegans*, named *nmur-1* to *nmur-3*. Boxes represent exons and lines show intronic sequences. Bars indicate the position of deletions in loss-of-function alleles. **(c)** Overview of the gustatory associative learning paradigm. The intrinsic attraction to NaCl is reversed by conditioning worms with salt in the absence of food during a short-term (15 min) training period. Mock-conditioned animals, which have not been pre-exposed to salt, are used as control. After conditioning, NaCl chemotaxis behavior is measured by determining the distribution of worms on quadrant plates of which two opposing quadrants are supplemented with NaCl and a chemotaxis index is calculated. **(d)** Gustatory associative learning of mutants defective in NMUR signaling. A mutant of *osm-9*, a TRPV channel subunit required for gustatory learning, is used as positive control^46^. In this and all subsequent figure, data from mock-conditioned worms is depicted in blue, while red markers denote NaCl-conditioned worms.

The learning defect of *nmur-1* animals could be caused by a failure to associate salt with the lack of food, or may alternatively result from general defects in locomotion or salt chemotaxis behavior. To address this, we tested *nmur-1* mutants for their behavioral responses to salt in different training regimes (Supplementary figure 1). We first examined chemotaxis behavior of *nmur-1* animals for increasing NaCl concentrations and observed no difference compared to wild-type animals (Supplementary figure 1a-b). To determine if food cues contribute to the plasticity defect, we performed conditioning on agar plates instead of pre-exposing animals to salt in liquid (Supplementary figure 1c). Mutants of *nmur-1* showed normal salt chemotaxis behavior when salt was paired with food (Supplementary figure 1d). Mock-conditioned *nmur-1* animals that were not pre-exposed to salt also behaved like wild-type worms (Figure 1d and Supplementary figure 1d). These results suggest that salt sensing is not affected in *nmur-1* mutants and that NMUR-1 signaling promotes gustatory aversive learning.

### NMUR-1 regulates distinct navigational decisions for the experience-dependent modulation of salt chemotaxis behavior

*C. elegans* utilizes two main behavioral strategies to navigate NaCl gradients. First, untrained worms increase the duration of forward movement (runs) when salt concentrations increase locally, called biased random walk^47^. Second, they actively steer towards attractive NaCl concentrations, known as klinotaxis^48^. After aversive conditioning with salt in the absence of food, both strategies are reversed to mediate NaCl avoidance instead of salt attraction^48,49^. As specific neuronal substrates have been delineated for each strategy^48–51^, we asked whether *nmur-1* is specifically involved in one of these responses, or if it acts as a general regulator of learned salt avoidance and thus modulates both behavioral strategies.

To answer this question, we set up a tracking platform to monitor trajectories of individual worms and quantify locomotion parameters relative to a linear salt gradient on square chemotaxis plates (Figure 2a). We first examined how gustatory aversive learning in wild-type animals is manifested on these linear salt gradients by measuring the position of worms during a 30-minute time window immediately after training. Similar to the quadrant-based assay (Figure 1c), mock-conditioned wild-type animals were attracted to salt and had migrated up the gradient after ten minutes (Figure 2b and c). NaCl-conditioning in the absence of food significantly reduced salt attraction, as conditioned animals avoided high NaCl concentrations (Figure 2b and c). Besides the positions of animals on the gradient, we determined a chemotactic index by dividing the mean velocity of each individual’s trajectory relative to the gradient by its mean crawling speed^22^. As expected, mock-conditioned wild-type animals had a positive chemotactic index, meaning they are attracted to salt, whereas NaCl-conditioned animals showed a negative index due to salt avoidance (Figure 2d). We also calculated indices to describe the two behavioral strategies for salt chemotaxis. In a biased random walk index, we determined the fractional difference in relative run duration up or down the gradient of individual trajectories (Figure 2e). The klinotaxis index was calculated as the fractional difference in the relative probability of each animal to reorient up or down the gradient after each turn (Figure 2f). In line with former studies^22,48^, we found that both navigational strategies are modulated by aversive experience, because the biased random walk and klinotaxis indices of wild-type animals reversed after NaCl-conditioning (Figure 2e – f).

**Figure 2.**
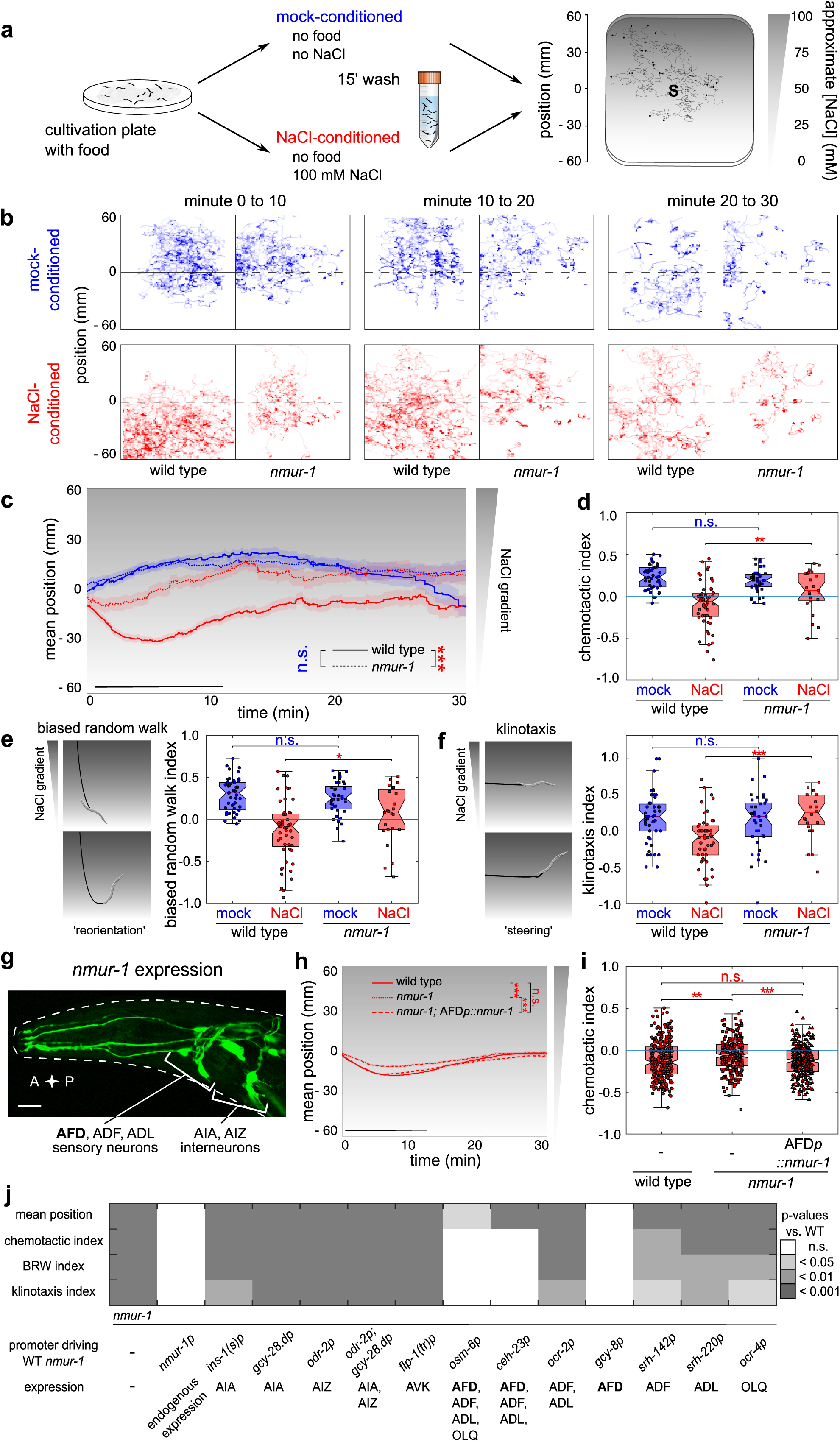
NMUR-1 signaling in AFD sensory neurons modulates distinct behavioral strategies underlying learned salt chemotaxis behavior (a) Schematic of behavioral tracking for gustatory learning. NaCl chemotaxis behavior is monitored on 0 to 100 mM linear NaCl gradients. After conditioning, worms are released in the middle of the gradient (S, source) and allowed to navigate for 30 minutes. Custom-written tracking software is used to extract positional data and determine behavioral indices for quantifying forward movement and turning behavior relative to the gradient. **(b)** Navigation trajectories of 25 - 50 wild type *C. elegans* and *nmur-1* mutant animals after mock- and NaCl-conditioning. Panels display the relative x-y residence on the linear NaCl gradients in 10-min intervals. **(c)** Average positions of wild type and *nmur-1* mutant worms on the linear NaCl gradient through time after mock- and NaCl-conditioning. n ≥ 25 animals per genotype. In this and all subsequent figures, black bars indicate the time interval used for statistical comparison of the mean position on the gradient, but also for calculating the chemotactic, biased random walk and klinotaxis indices in panels d-f. Shaded regions represent S.E.M. **(d)** The chemotactic index quantifies the preferential migration vector of each individual worm. n ≥ 25 animals per genotype. **(e)** Quantification of biased random walk, assessing the relative duration of forward runs when navigating up the salt gradient. n ≥ 25 animals per genotype. **(f)** Quantification of klinotaxis behavior. The klinotaxis index reflects the behavioral strategy by which worms actively steer towards attractive NaCl concentrations. n ≥ 25 animals per genotype. **(g)** Transgenic adult hermaphrodite *C. elegans* expressing a bicistronic GFP construct including the promoter, cDNA and 3’UTR of the *nmur-1* gene (*nmur-1p::nmur-1::SL2::gfp*). Scale bar, 35 µm. **(h)** Mean position on the NaCl-gradient and **(j)** chemotactic index of NaCl-conditioned *nmur-1* mutant worms in which *nmur-1* expression is specifically restored in the AFD sensory neurons by expressing *nmur-1* cDNA under the control of the *gcy-8* promoter^53^. n ≥ 284 animals per genotype. **(j)** Summary statistics of endogenous and cell-specific rescue experiments of learned salt aversion. For each of four behavioral parameters, significances are relative to NaCl-conditioned wild-type animals tested on the same day. Raw data are depicted on Supplementary Fig. 3 and 4, for cell-specific rescue in the interneurons or sensory neurons, respectively. n ≥ 50 animals per genotype.

Using this behavioral tracking platform, we further examined the defect in learned salt avoidance of *nmur-1* mutants. Similar to the quadrant-based assay (Figure 1d), mock-conditioned *nmur-1* animals displayed normal salt chemotaxis behavior (Figure 2c-f), while NaCl-conditioned mutants showed less aversion for high NaCl concentrations than wild-type animals (Figure 2b-d). Interestingly, we observed significant differences in biased random walk and klinotaxis of *nmur-1* mutants after aversive conditioning, suggesting that NMUR-1 signaling is involved in the experience-dependent modulation of both the length of runs and the reorientation on the NaCl gradient (Figure 2f and g).

To confirm the role of *nmur-1* in gustatory aversive learning, we tested whether restoring the expression of *nmur-1* rescued the mutant’s phenotypes. Therefore, we expressed wild-type copies of the receptor under the control of its endogenous promoter (Figure 2g), which recapitulated the reported expression pattern for *nmur-1* in approximately 15 sensory neurons (ADF, ADL and AFD) and interneurons (AIA, AIZ, among others)^52^. This transgene fully rescued salt chemotaxis behavior of *nmur-1* mutants after aversive conditioning, including defects in the modulation of klinotaxis and biased random walk (Figure 2j and Supplementary figure 2a for raw data). Next, we asked in which cell(s) NMUR-1 acts to potentiate salt aversive learning by probing each *nmur-1* expressing neuron in cell-specific rescue experiments (Figure 2j and Supplementary figures 2-3 for raw data). Re-introducing the wild-type *nmur-1* cDNA in AFD sensory neurons using various promoters, including the AFD-specific *gcy-8* promoter^53^ fully restored learned salt aversion, whereas cell-specific expression in other *nmur-1* expressing sensory neurons and interneurons did not restore the mutant phenotype (Figure 2h, i, j and Supplementary figure 3a, b and d). Taken together, the NMU receptor ortholog NMUR-1 is required in the AFD sensory neurons to coordinately modulate distinct navigational decisions for establishing learned salt avoidance.

### NMU-like neuropeptides of the *C. elegans* CAPA-1/NLP-44 precursor are ligands of NMUR-1

NMUR-1 is an orphan GPCR, meaning that so far no ligands have been identified for this receptor. To further characterize the role of NMUR-1 in gustatory aversive learning, we set out to identify its ligand(s) using reverse pharmacology. Therefore, we expressed NMUR-1 in Chinese Hamster Ovary (CHO) cells that were challenged with a synthetic library of over 300 *C. elegans* peptides, and we monitored GPCR activation by a calcium reporter assay (Figure 3a). Only neuropeptides encoded by the *capa-1/nlp-44* gene elicited a calcium response in NMUR-1 expressing cells. Cells transfected with an empty control vector showed no response (Figure 3b).

**Figure 3.**
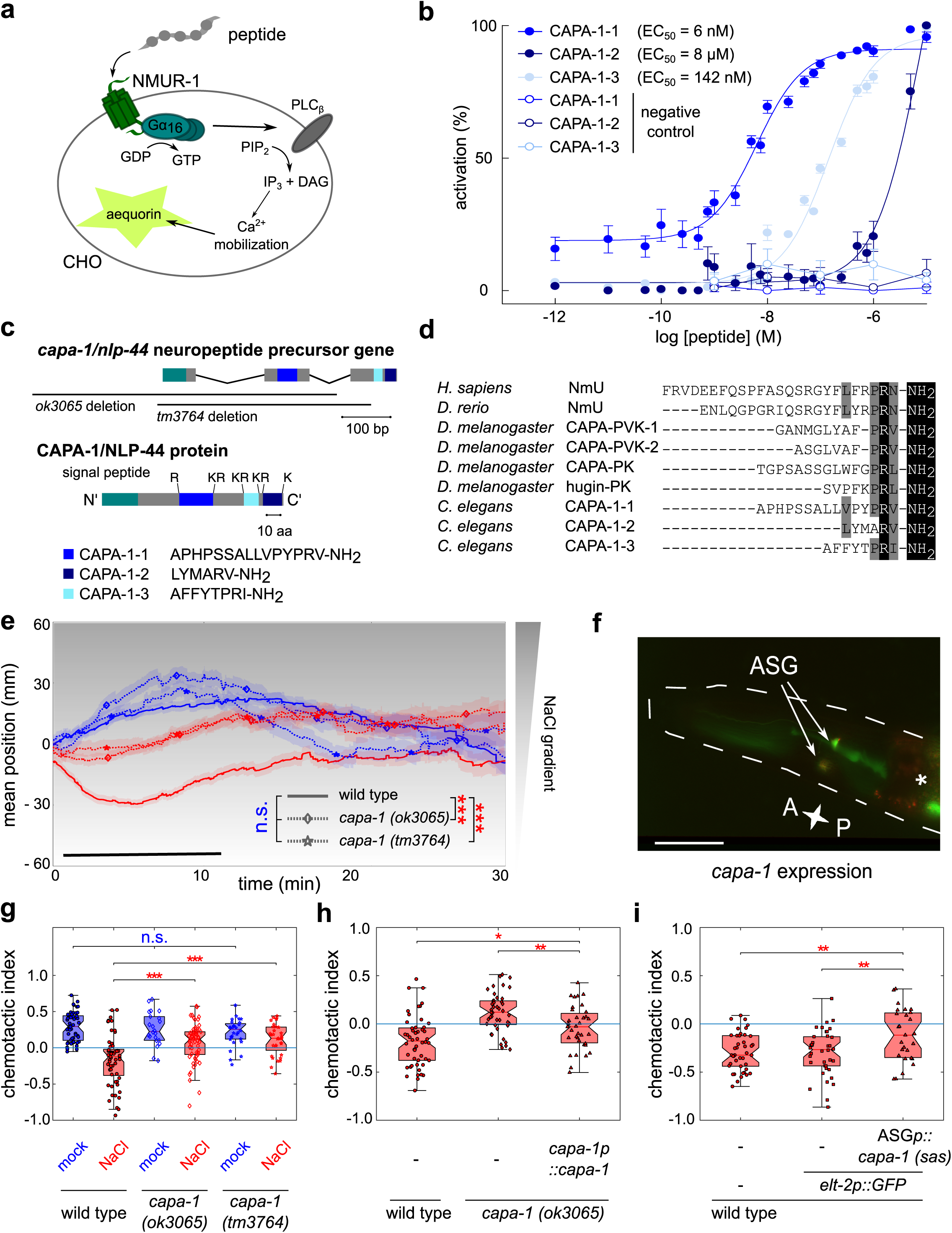
CAPA-1 NMU-like neuropeptides signal from ASG chemosensory neurons to promote gustatory learning through NMUR-1 (a) Luminescence-based calcium mobilization assay for measuring GPCR activation. NMUR-1 is expressed in CHO cells that stably co-express the promiscuous human Gα_16_ subunit, which couples receptor activation to calcium release from intracellular storage sites. Intracellular calcium levels are monitored using the calcium indicator aequorin. **(b)** Calcium responses of CHO cells expressing NMUR-1 are shown relative (%) to the highest value (100% activation) after normalization to the total calcium response. Cells transfected with an empty pcDNA3.1 vector are used as a control. Error bars show S.E.M (n ≥ 6). **(c)** The *capa-1/nlp-44* gene encodes a neuropeptide precursor that harbors three predicted peptide sequences (CAPA-1-1, -2, and -3), which are flanked by mono- and dibasic cleaving sites. Black bars indicate the positions of deletions in *capa-1/nlp-44* mutant alleles used in this study. **(d)** Sequence comparison of *C. elegans* CAPA-1 neuropeptides and NmU neuropeptides from *H. sapiens*, *D. rerio*, and *D. melanogaster*. Conserved C-terminal features are highlighted in black. Residues with similar physicochemical properties are colored gray. **(e)** Mean position on the gradient through time and **(g)** chemotactic indices of two independent deletion mutants of the *capa-1/nlp-44* neuropeptide precursor gene after mock- and NaCl-conditioning. The corresponding biased random walk and klinotaxis indices are shown on Supplementary Fig. 5a. n ≥ 25 animals per genotype. **(f)** Co-localization of a bicistronic GFP construct harboring the promoter, genomic DNA and 3’UTR of the *capa-1* gene *(capa-1p::capa-1::SL2::gfp*) with mCherry expression from the ASG-specific *gcy-15* promotor^55^. The asterisk marks fluorescence in the intestine from the *elt-2p::gfp* co-injection marker. Scale bar, 35 µm. **(h)** Chemotactic behavior of NaCl-conditioned *capa-1 (ok3065)* animals in which wild type copies of the *capa-1* gene are reintroduced under the control of its promoter sequence (*capa-1p::capa-1::SL2::gfp*). The corresponding mean population position on the gradient, biased random walk and klinotaxis indices are shown on Supplementary Fig. 5b. n ≥ 37 animals per genotype. **(i)** Chemotactic behavior of NaCl-conditioned animals after ASG-specific RNAi-mediated knockdown of *capa-1*, achieved by expressing *sense* and *anti-sense capa-1* sequences under the control of the ASG-specific *ops-1* promoter^60^. A strain carrying only the *elt-2p*::GFP co-injection marker is used as a control to exclude potential effects of the transgene selection marker. The corresponding mean population position on the gradient, biased random walk and klinotaxis indices are shown on Supplementary Fig. 5c. n ≥ 27 animals per genotype.

*C. elegans* NLP-44 is a neuropeptide precursor that belongs to the NMU family^32,35^. It encodes three predicted peptides with a C-terminal YXPR(I/V)-NH_2_ motif, which resembles the bioactive PRX-NH_2_ motifs of *Drosophila* and mammalian NMU-like ligands^54^ (Figure 3d). The *Drosophila* NMU signaling system consists of pyrokinin peptides encoded by the *capability* and *hugin* genes. Due to the sequence similarity of *nlp-44* with the insect *capability* gene*, C. elegans nlp-44* has been named *capa-1*^35^. We determined the potency of synthetic CAPA-1 peptides to activate NMUR-1 by measuring GPCR activation for increasing peptide concentrations. CAPA-1-1 activated NMUR-1 with a half maximal effective concentration (EC_50_) of 6 nM, while CAPA-1-2 and CAPA-1-3 showed markedly higher EC_50_ values of 8 µM and 142 nM, respectively (Figure 3b).

### NMU-like CAPA-1 neuropeptides signal from ASG sensory neurons to promote gustatory aversive learning

Because CAPA-1 peptides activate NMUR-1 *in vitro*, we hypothesized that these neuropeptides modulate gustatory aversive learning through NMUR-1 signaling. For this purpose, we examined gustatory learning in two *capa-1* mutants with independent loss-of-function alleles (Figure 3c). As expected, we found that both *capa-1* mutants display a learning defect, which mimics that of *nmur-1* mutants (Figure 3e and g). Impaired CAPA-1 signaling, however, has no effect on NaCl chemotaxis behavior of mock-conditioned animals. Like *nmur-1* mutants, worms lacking CAPA-1 signaling show defects in the modulation of biased random walk and klinotaxis after NaCl-conditioning (Supplementary figure 4a). The reduced salt avoidance of *capa-1* mutants does not result from general defects in locomotion or salt chemotaxis, because loss of *capa-1* function did not cause a detectable change in behavioral responses to a range of NaCl concentrations and mutants of *capa-1* behaved like wild-type animals after pairing salt with the presence of food (Supplementary figure 5a and b). These results suggest that NMU-like CAPA-1 neuropeptides are also required for gustatory aversive learning.

Using a bicistronic GFP reporter transgene, we found that *capa-1* is expressed in a single pair of neurons (Figure 3f), which is in accordance with previous whole-mount immunostaining using an antiserum that recognizes CAPA-1-derived neuropeptides^35^. Based on the cellular position and morphology, we identified these cells as the ASG neurons. Their cellular identity was further confirmed by co-localization of the GFP signal with an mCherry protein expressed from the *gcy-15* promoter (Figure 4f) that is known to drive expression in ASG only^55^. ASG is a chemosensory neuron pair that is involved in detecting food and water-soluble attractants such as NaCl^56–58^. Restoring *capa-1* expression in ASG under the control of its promoter rescued the learning defect of *capa-1* mutants (Figure 3h and Supplementary figure 4b). To further validate ASG as the source of CAPA-1 release in gustatory learning, we knocked down *capa-1* expression specifically in these cells by expressing sense and antisense sequences of *capa-1* under the control of an ASG-specific (*ops-1*) promoter^59,60^. ASG-specific knockdown of *capa-1* resulted in a gustatory learning defect, because both the migration on the salt gradient and the behavioral indices differed significantly between animals with knockdown of *capa-1* and animals that don’t express the RNAi transgene (Figure 3i and Supplementary figure 4c). CAPA-1 signaling from ASG neurons thus modulates salt chemotaxis behavior according to previous experience.

**Figure 4.**
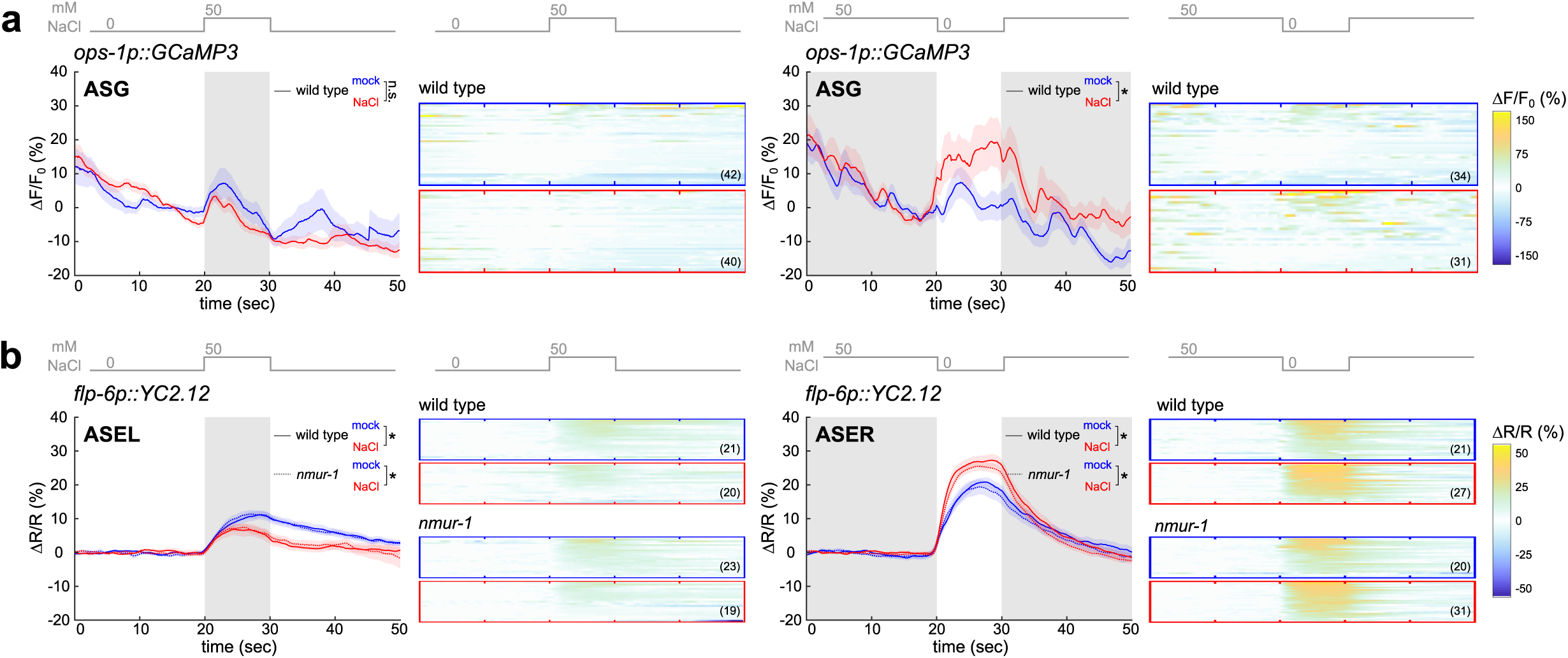
ASG and ASE salt-induced activity are shaped by NaCl-conditioning (a) NaCl-induced calcium responses in ASG neurons of mock- and NaCl-conditioned animals. Wild type animals expressing the GCaMP3 calcium indicator in ASG are challenged to either 0 to 50 mM NaCl up-step (left panel) or 50 to 0 mM NaCl down-step (right panel) after pre-exposure to 0 mM NaCl (mock-conditioned, blue traces) or 100 mM NaCl (NaCl-conditioned, red traces) for 15 min without food in a microfluidic chip. Calcium dynamics are relative to prestimulus control values, ΔF/F_0_. **(b)** NaCl-induced calcium responses in ASE neurons of mock- and NaCl-conditioned wild-type or *nmur-1* mutant animals. Worms expressing the YC2.12 calcium indicator in ASE are challenged to either 0 to 50 mM NaCl up-step (left panel) or 50 to 0 mM NaCl down-step (right panel) after pre-exposure to 0 mM NaCl (mock-conditioned, blue traces) or 100 mM NaCl (NaCl-conditioned, red traces) for 15 min without food in a microfluidic chip. Traces indicate the average percent change in the ratio of the distinct wavelengths of the YC2.12 indicator.

We previously reported that one of the CAPA-1 peptides, CAPA-1-3, activates NMUR-2, a second neuromedin receptor ortholog^35^. Although *nmur-2* single mutants were not defective in gustatory aversive learning (Figure 1d and Supplementary figure 6), NMUR-1 signaling may mask a role of this receptor in the NMU-dependent modulation of salt chemotaxis behavior. We therefore compared learning in *nmur-1* and *nmur-2* single mutants with a double mutant of the two receptors. The defect of double mutant animals closely resembled that of the *nmur-1* single mutant (Supplementary figure 6), suggesting that only NMUR-1 mediates CAPA-1 signaling in gustatory learning.

### CAPA-1/NMUR-1 signaling is not involved in food searching behaviors

Previous work indicates that the ASG neurons, which express CAPA-1, regulate foraging behaviors^61^. NMU-like peptides are potent regulators of feeding in insects and mammals^62,63^, and the *C. elegans* receptor NMUR-1 has also been shown to modulate lifespan depending on the type of bacterial diet^52,64^. These observations prompted us to investigate if the CAPA-1/NMUR-1 pathway is involved in signaling the environmental food context and if this function could underlie its effect on gustatory aversive learning.

Loss of NMUR-1 has been shown to extend lifespan only in worms grown on B-type *E. coli* strains such as OP50, while it has no effect on K-12 type bacterial strains like HT115. We therefore asked if the effect of NMUR-1 signaling on gustatory aversive learning depends on the bacterial diet, by assaying gustatory plasticity of worms cultured on HT115 *E. coli* (Supplementary figure 7). Mutants of *capa-1* and *nmur-1* grown on HT115 showed learning defects similar to worms cultured on OP50 (Figure 2c-f; Figure 3e and g and Supplementary figure 4a). Impaired NMU-like signaling hence disrupts gustatory learning independent of the bacterial diet.

We also tested whether CAPA-1/NMUR-1 signaling is involved in ASG-dependent foraging behaviors. Upon removal from their *E. coli* food source*, C. elegans* display an intensive local search behavior with frequent turning and reorientations, which are reduced in ASG-ablated animals^65,66^. When we removed mutants for *capa-1* or *nmur-1* from their OP50 or HT115 food source, we observed no difference in local search behavior compared to wild-type *C. elegans* (Supplementary figure 8), indicating that NMU-like signaling is not involved in this behavioral state.

Besides local search behavior off food, ASG neurons have been shown to modulate foraging when worms are feeding on *E. coli* bacterial lawns. Under these conditions*, C. elegans* spontaneously alternate between active exploratory behavior, called roaming, and a more sedentary behavior, called dwelling^67,68^. ASG signaling is involved in regulating the switch between these behavioral states and promotes roaming^61^. To determine if *capa-1* and *nmur-1* regulate foraging on food, we quantified the relative time that the mutants spent roaming or dwelling, and calculated their mean speed. Our results show that foraging is not affected in *capa-1* or *nmur-1* mutants feeding on OP50 or HT115 bacteria (Supplementary figure 8g-j). Taken together, these findings indicate that CAPA-1 signaling is not generally involved in food sensing, because *capa-1* and *nmur-1* mutants display normal food searching behaviors.

### Aversive NaCl-conditioning shapes calcium responses of *capa-1* expressing ASG neurons to salt stimuli

Since CAPA-1 neuropeptides do not seem to be required for sensing salt or food conditions, we hypothesized that CAPA-1 signaling from ASG neurons modulates the neural processing of salt cues after aversive conditioning. Consistent with this model, laser ablation and calcium imaging studies in untrained wild-type animals suggest only a minor role for ASG neurons in salt sensing^58,69,70^. However, these neurons have been implicated in modulating behavioral responses to salt. For example, hypoxic stress increases attraction to NaCl through ASG signaling^71^.

The potential of ASG to modulate gustatory responses under certain conditions prompted us to compare ASG calcium activity in response to NaCl stimuli in mock- and NaCl-conditioned wild-type *C. elegans*. To this end, we used transgenic worms expressing the calcium indicator GCaMP3 in ASG neurons, and challenged individual worms with salt stimuli in a microfluidic chip^69,72^. Animals were conditioned with protocols similar to those used for behavioral learning assays, i.e. we first pre-exposed worms to a buffer with (NaCl-conditioned) or without (mock-conditioned) NaCl for 15 minutes before testing ASG activity. We then measured ASG calcium responses in response to either up- or downshifts in NaCl concentrations^73^. ASG showed small calcium responses to NaCl in both mock- and NaCl-conditioned animals. However, the response to NaCl downshifts was higher in worms pre-exposed to salt in the absence of food. After aversive conditioning, ASG responded significantly stronger to NaCl removal in comparison to ASG responses in mock-conditioned worms (Figure 4a). This result suggests that CAPA-1 neurons respond differently to salt stimuli after aversive conditioning and may facilitate learning through modulation of the salt sensing circuit.

Next, we asked if NMU signaling adapts salt-evoked activity of the primary salt sensor, ASE. This left-right pair of neurons (ASEL and ASER), exhibits functional asymmetry: ASEL and ASER are activated by an increase and decrease in NaCl concentrations, respectively^70,74^. Consistent with this, mock-conditioned wild-type worms expressing the ratiometric YC2.12 indicator in both ASEL and ASER showed robust ASEL responses upon NaCl upshifts, while a downshift in NaCl concentration reliably evoked an increase in calcium in the ASER neuron (Figure 4b). Following NaCl-conditioning, ASEL responses to a NaCl upshift were significantly reduced and ASER responses to NaCl downshifts were sensitized, which is in agreement with previous studies^73,75^. ASE calcium responses to salt in mock- and NaCl-conditioned animals were not affected in *nmur-1* mutants, indicating that NMU signaling is not required for the plasticity of ASE responses during aversive learning.

### CAPA-1 neurons are required for the retrieval, but not the acquisition, of learned salt avoidance

To gain further insight into the mechanisms by which NMU-like neuropeptides regulate aversive learning, we set out to uncover the temporal requirement of CAPA-1 neurons in the learning circuit. Therefore, we first investigated the effect of chemically silencing ASG by expressing tetanus toxin light chain (TeTx) specifically in these neurons. TeTx disrupts chemical signaling as it impedes the release of synaptic vesicles and neuropeptide-containing dense core vesicles^76^. As expected from the role of *capa-1* in gustatory plasticity, ASG-specific expression of TeTx impaired aversive learning, whereas it did not affect NaCl chemotaxis of mock-conditioned animals (Figure 5a-d).

**Figure 5.**
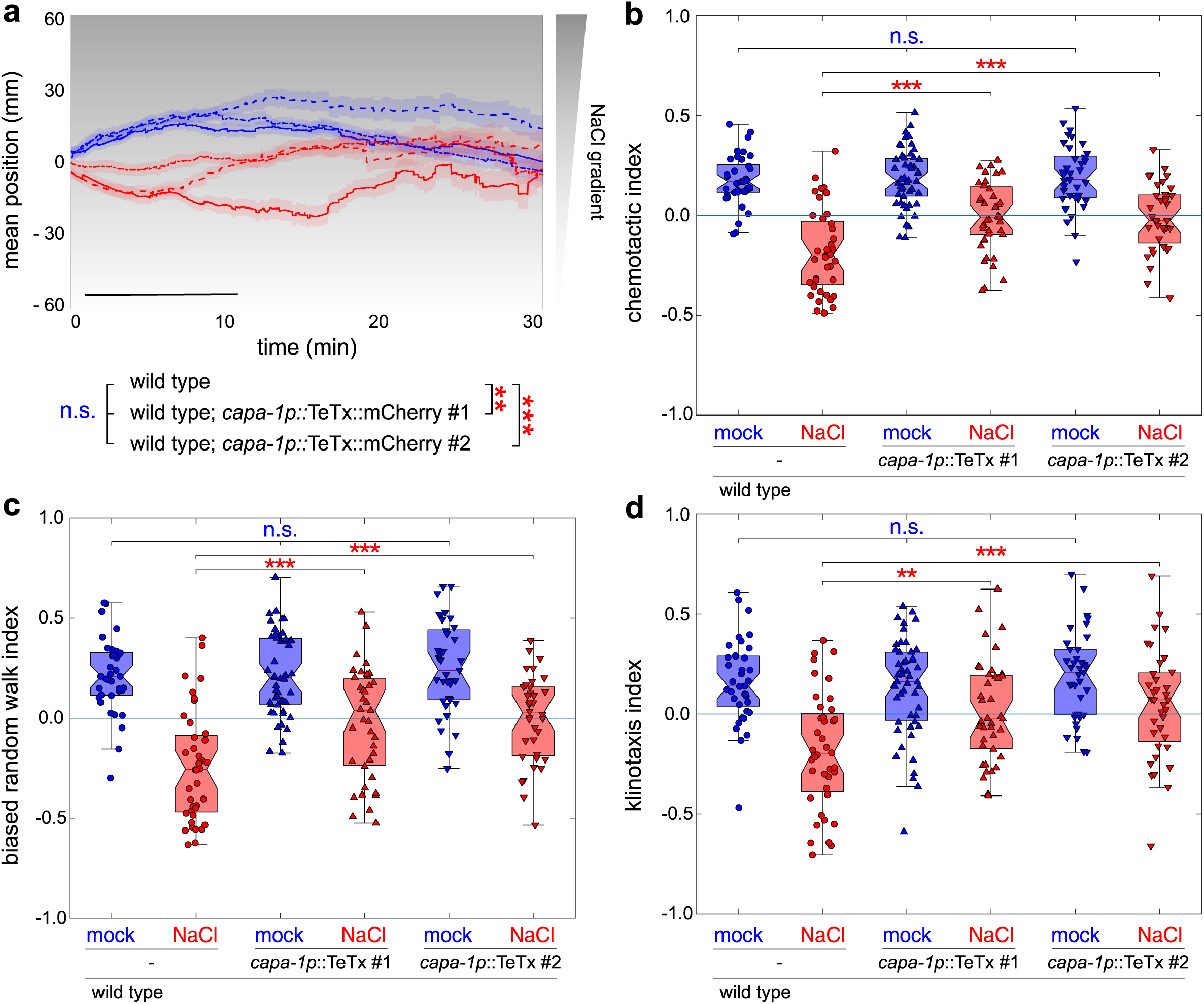
ASG chemical signaling is required for gustatory associative learning (a-d) Mock- and NaCl-conditioned behavior of two independent transgenic animals expressing the tetanus toxin light chain (TeTx) in ASG, which blocks synaptic release (*capa-1p::TeTx::mCherry)*. **(a)** Mean population position on a 0 – 100 mM NaCl gradient after mock- (blue traces) or NaCl-conditioning (red traces). Shaded regions represent S.E.M. Black bar indicates the time interval for statistical comparison of the mean position on the gradient, but also for calculating the chemotactic **(b)**, biased random walk **(c)** and klinotaxis **(d)** indices.

To further determine when ASG signaling is required for gustatory learning, we optogenetically silenced these neurons during NaCl-conditioning or during the immediate recall of learned avoidance behavior when worms are navigating on a NaCl gradient. For this purpose, we expressed the outward-directed proton pump Arch under the control of the *capa-1* promoter^77,78^ (Figure 6a). Upon adding the essential cofactor for opsin activity (*all-trans* retinal, ATR), Arch can be activated by illuminating transgenic worms with green-red light. The subsequent membrane hyperpolarization silences the host neuron. We varied the timing of illumination so that CAPA-1 neurons were silenced either during the acquisition of salt aversion or during retrieval of the learned behavior (Figure 6b). Silencing CAPA-1 neurons during the acquisition phase had little effect on learned salt avoidance (Figure 6c-f). In contrast, silencing during the retrieval phase significantly attenuated salt avoidance (Figure 6c-f). Optogenetically inhibiting CAPA-1 neurons had no effect on general locomotion in the absence of a NaCl gradient (Supplementary figure 9). These results show that CAPA-1 neurons are required for the retrieval of learned salt avoidance, but not for the acquisition of the conditioned response.

**Figure 6.**
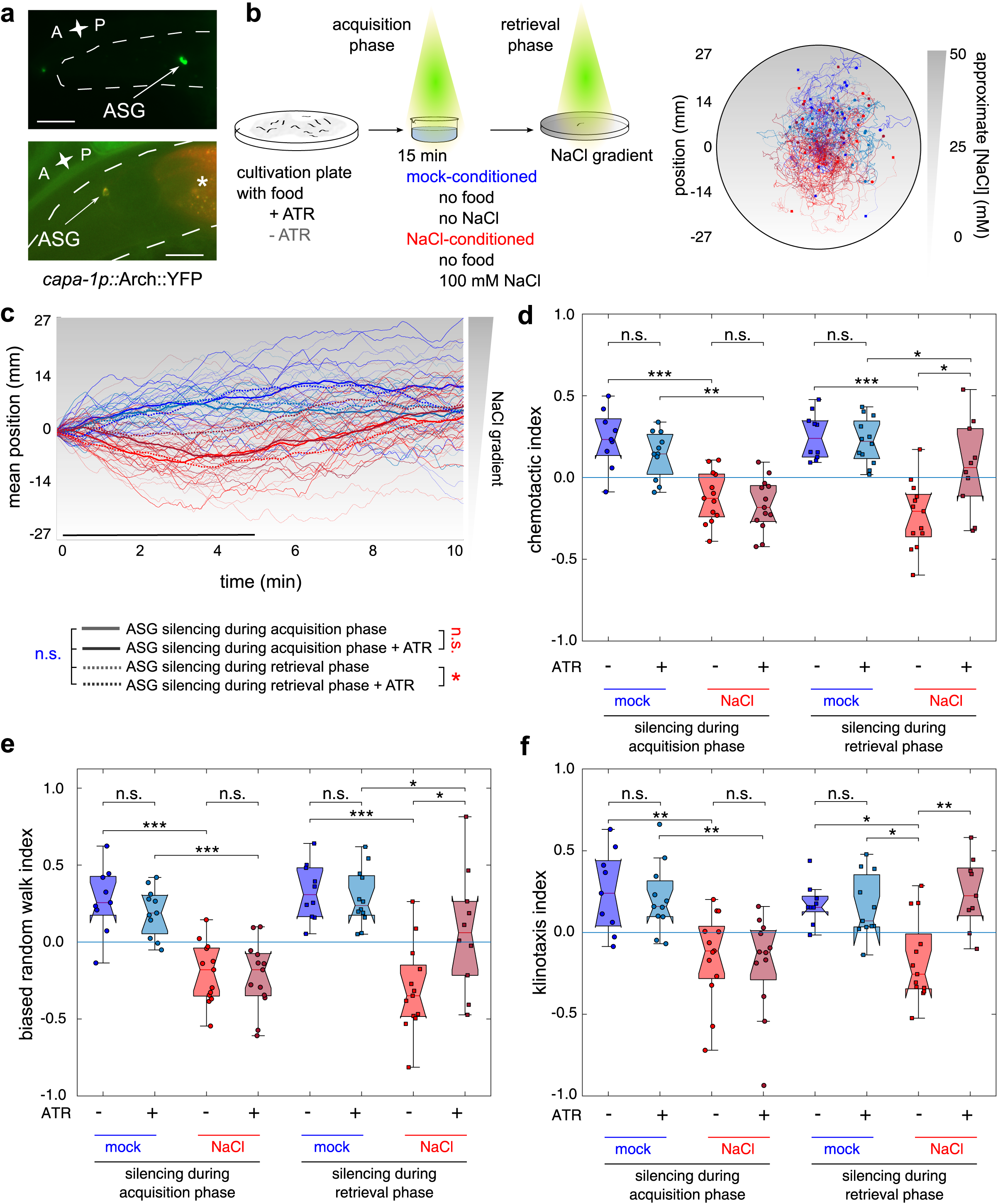
Optogenetic silencing of CAPA-1 neurons during retrieval of learned salt aversion attenuates NaCl-conditioned behavior (a) The inhibitory opsin Arch is expressed under control of the *capa-1* promoter (top panel), and co-localization of the Arch::YFP signal assessed with mCherry expressed from the ASG-specific *gcy-15* promoter (bottom panel). **(b)** Individual worms are illuminated with yellow-green light for silencing CAPA-1 neurons during conditioning (acquisition phase) or during the NaCl chemotaxis assay when worms are navigating a linear NaCl gradient (retrieval phase). Arch requires the cofactor *all-trans* retinal (ATR) to be supplemented to the worm culture. Transgenic *C. elegans* not fed ATR-supplemented food serve as a negative control. After conditioning, worms are put in the middle of a chemotaxis plate with a linear NaCl gradient (0 to 50 mM) and are kept in the field-of-view using an automated xy-stage. Right panel shows navigation trajectories of ≥ 10 worms per condition. **(c)** Positions of individual worms on the NaCl gradient through time. Thick lines represent mean positions. Black bar indicates the time interval for calculating chemotactic, biased random walk and klinotaxis indices in panels d-f. **(d-f)** CAPA-1 neurons are not required for NaCl chemotaxis behavior of mock-conditioned animals, because ATR-fed animals display normal chemotactic, biased random walk and klinotaxis behavioral indices. Optogenetic silencing of CAPA-1 neurons after NaCl-conditioning during the retrieval phase disrupts the learned salt avoidance response, while silencing during acquisition has no effect.

## Discussion

Although neuropeptides are increasingly recognized as modulators of learning and memory, it is not well understood how they adapt behavioral choices and when specific neuromodulators operate in learning circuits. In this study we demonstrate that neuropeptides of the NMU family promote gustatory aversive learning in *C. elegans* by modulating distinct navigational decisions according to previous experience. We show that NMU neurons are specifically required for the retrieval of learned avoidance behavior. Since NMU neuropeptides are highly conserved in bilaterian animals, including humans, these findings lay a foundation for further research on the involvement of this ancient neuropeptide family in other learning circuits.

In *C. elegans* NMU-like neuropeptides, encoded by the *capa-1/nlp-44* gene, are expressed in a single pair of chemosensory neurons, the ASG cells, which promote gustatory aversive learning through CAPA-1 signaling. To determine when signaling from CAPA-1 neurons is required for learning, we used optogenetics to temporally silence these cells during the conditioning phase or during retrieval of the conditioned response immediately after training. Only the latter affected the experience-dependent adaptation of NaCl chemotaxis. CAPA-1 neurons thus control the expression of avoidance behavior, but are not required for acquisition of this response. Evidence for such a temporally defined role of neuropeptide signaling in learning circuits also emanates from *Drosophila* studies on neuropeptide F and insulin-like peptides, which are large neuropeptides of over 100 amino acids that act via GPCR and receptor tyrosine kinase signaling, respectively^79–81^. Of notice, one *C. elegans* study characterized the memory phase of insulin receptor signaling in an olfactory learning circuit.

Using a conditional *daf-2* allele, the authors showed that insulin receptor signaling in the olfactory AWC neurons is only partially involved in memory acquisition, but necessary during memory retrieval^82^. By manipulating neuropeptide signaling at the level of peptidergic neurons, our results indicate that small neuropeptide messengers of the NMU family specifically control memory retrieval. A defined temporal role of neuropeptides in learning circuits may be a general feature of neuropeptide-mediated memory.

How does CAPA-1 signaling regulate the expression of learned salt avoidance behavior? We identified the NMU receptor ortholog NMUR-1 as a CAPA-1 receptor that is required for gustatory aversive learning. NMUR-1 is present in several sensory neurons and interneurons and has previously been implicated in the food-dependent modulation of lifespan^52,64^. However, the learning defects of *capa-1* and *nmur-1* mutant animals are not affected by the worms’ bacterial diet and CAPA-1 signaling mutants display normal foraging, salt-sensing and locomotion behaviors. Therefore, a general function in sensing salt or the environmental food context does not seem to underlie the effect of CAPA-1 peptides on aversive learning. Our calcium imaging data suggest a model in which aversive conditioning recruits CAPA-1 neurons to modulate the gustatory circuit. Aversive conditioning sensitizes the response of CAPA-1 neurons to NaCl downshifts, which suggests that ASG neurons contribute to the experience-dependent adaptation of salt chemotaxis behavior when worms are exposed to the aversive condition of salt without food. This function is reminiscent of a previous report showing ASG to be recruited to a circuit for processing NaCl cues under another type of stressful condition, hypoxia^71^. After prolonged exposure to hypoxic stress, ASG neurons enhance NaCl attraction through increased serotonin production. Whether CAPA-1 neuropeptides are involved in this process is not known.

At the behavioral level, CAPA-1/NMUR-1 signaling facilitates the adaptation of two elementary behavioral strategies for NaCl chemotaxis, i.e. biased random walk and klinotaxis. NMU-like signaling thus acts as a central regulator of different decision-making tasks for expressing learned avoidance behavior, rather than specifically affecting a neural substructure of the behavioral output. Conform to previous work, we found gustatory aversive conditioning to adapt the activity of the primary salt-sensor ASE that mediates the attractive drive towards NaCl^73,75^. This happens in an NMUR-1-independent manner, suggesting that NMU signaling does not indirectly attenuate the neural pathway for NaCl attraction. By contrast, we found that CAPA-1 facilitates salt learning by acting on NMUR-1 in AFD sensory neurons, which play a well-characterized role in various sensory modalities such as the sensation of thermal, magnetic and CO_2_ cues^83–85^. Interestingly, AFD has recently been shown to respond to NaCl fluctuations^86^ and wires to an interconnected layer of interneurons involved in transforming salt cues into appropriate behavioral responses^28^. As this layer encompasses neuronal elements driving biased random walk and klinotaxis strategies^48–51^, AFD signaling may acutely impinge on the gustatory circuit downstream of the primary salt sensor ASE to modulate navigational decisions and change the NaCl valence from attractive to aversive. Taken together, these findings argue for a model in which the CAPA-1 neurons have no primary role in innate salt attraction, but are engaged under stressful or aversive conditions and recruit another sensory neuron within the circuit for salt chemotaxis. This neuropeptide-driven modulation may increase the sensory acuity of the animal when faced with an aversive environment.

Would similar neuropeptidergic systems be involved in other learning circuits? The NMU signaling system is ancestral to bilaterian animals^32–34^ and has been shown to regulate energy homeostasis in vertebrates and insects^62,87–91^. A small number of studies have postulated a role for NMU signaling in memory circuits in mammals, based on the observation that intranasal administration or injection of NMU peptides into the brain affects memory and reward-related behaviors^39–43^. For one type of memory, this effect of NMU peptides has been delineated to a neuroprotective function rather than the modulation of sensory processing^39,40^. In *Drosophila* a subset of neurons expressing the NMU precursor gene *hugin* is part of a sensory pathway that relays between gustatory neurons and higher brain centers. This neural subset transduces aversive gustatory cues, such as bitter tastes, resulting in a brake on feeding through avoidance behavior and decreased pharyngeal pumping^38,89,92,93^. In this context, *hugin* may play a role similar to *capa-1* in *C. elegans*, signaling aversive feeding conditions and stimulating motor programs that allow the animal to migrate to more favorable food environments. Since both the *hugin* gene and the *capa* receptor gene (*capaR*) are expressed in the mushroom bodies, the fruit fly’s brain centers that play an important role in sensory integration and memory formation^36^, the *Drosophila* NMU homolog might be involved in learning as well. Similarly, the NMU neuropeptides and the neuromedin receptor NMUR2 are expressed in discrete regions of the mammalian brain that are instrumental for memory and processing of aversive experiences, including the amygdala, hippocampus and cerebral cortex^37,94,95^.

Taken together, we have uncovered a neural mechanism by which the retrieval of learned avoidance is regulated by the evolutionarily conserved NMU neuropeptide pathway. Given that the NMU system is conserved across bilaterians and that NMU specifically regulates retrieval of learned avoidance, our study encourages further research into the memory functions of this neuropeptide system in other animals and in the context of aversive learning disorders.

## Acknowledgements

We thank members of the Schoofs and Temmerman labs for their thoughtful comments. The authors would like to thank F. J. Naranjo-Galindo for technical assistance. We thank the *Caenorhabditis* Genetics Center, supported by the NIH Office of Research Infrastructure Programs Grant P40 OD010440, the Mitani lab and the National Bioresource Project of Japan, the Alcedo lab (Wayne State University), the Bargmann lab (Rockefeller University), the Jorgensen lab (University of Utah), the Murayama lab (OIST Graduate University),S. Husson and the Gottschalk lab (Johann Wolfgang Goethe-Universität Frankfurt) and the Schafer lab (MRC Laboratory of Molecular Biology) for providing *C. elegans* strains and reagents. J.W., K.P., C.B., I.R. and I.B. are fellows of the Research Foundation - Flanders (FWO). This research was supported by European Research Council Grant 340318.

## Author Contributions

J.W., P.V.d.A., K.P., and I.B. conceived and designed experiments. J.W., C.B. and E.V. performed molecular genetics and worm transgenesis. J.W. performed all behavioral and optogenetic experiments. J.W. and P.V.d.A. conducted calcium imaging. I. R. and J. L. assisted with microfluidic chip construction. R. J. advised the construction of the imaging platform. J.W., P.V.d.A., K.P. and I.B. analyzed and interpreted results. J.W., L.S., and I.B. wrote the manuscript.

## Methods

Further information and requests for resources and reagents should be directed to and will be fulfilled by Liliane Schoofs and Isabel Beets.

### *C. elegans* strains

*C. elegans* was cultured using standard methods at 20°C on agar with Nematode Growth Medium (NGM) seeded with OP50 or HT115 *E. coli* bacteria as a food source. Wild-type worms are of the Bristol variety N2. N2 wild type and RB1288 *nmur-1 (ok1387)*, RB2526 *nmur-2 (ok3502)*, VC1974 *nmur-3 (ok2295)*, RB2263 *capa-1 (ok3065)*, CX10 *osm-9 (ky10)* mutants were acquired from the *Caenorhabditis* Genetics Center at the University of Minnesota. *capa-1 (tm3764)* mutants were provided by the National Bioresource Project of Japan. Mutant genotypes were confirmed using PCR and backcrossed to the common wild-type strain (N2) at least 6 times to remove unlinked mutations prior to analysis. All mutant and transgenic strains used in this study are listed in the additional resource table.

### Phylogenetic tree

To determine the relationship between NMU-like receptors from *Caenorhabditis elegans*, *Lottia gigantea, Drosophila melanogaster, Danio rerio* and *Homo sapiens*, a phylogenetic tree was generated by the maximum likelihood method using the phylogeny pipeline set in ‘advanced’ mode with the number of bootstraps set to 100 (www.phylogeny.fr)^1,2^. The sequences used to construct the tree are: *Homo sapiens* NMUR1 (AAH51914.1) and NMUR2 (EAW61653.1); *Danio rerio* Nmur-1a (ENSDARG00000060884), Nmur-1b (ENSDARG00000003944) and Nmur-2 (ENSDARG00000022570)^3^; *Drosophila melanogaster* capa receptor (CG14575), pyrokinin-1 receptor (A197.2), pyrokinin-2 receptor 1 and 2 (CG8784 and CG8795); *Lottia gigantea* NMUR (ULotgi1_91538); *Caenorhabditis elegans* NMUR-1 (NP_509515.3), NMUR-2 (CCD67735.1) and NMUR-3 (NP_001024525.1). Human (NP_003292.1), *D. rerio* (XP_687246.4, NP_001108160.1 and NP_001037811.1), *L. gigantean* (Lotgi1_122499) and *C. elegans* (Q95YD7) sequences of the thyrotropin-releasing hormone receptor family were chosen as an outgroup to root the tree.

### Molecular biology

All oligonucleotide primers used in this study are listed in the additional resource table.

For heterologous expression of NMUR-1 in Chinese hamster ovary (CHO) cells, *nmur-1* cDNA was amplified by PCR from cDNA of mixed-stage wild-type *C. elegans* and directionally cloned into the eukaryotic expression vector pcDNA3.1 (Invitrogen).

GFP reporter and rescue constructs were generated from a modified pSM vector carrying a GFP reporter sequence preceded by an SL2 trans-splicing sequence (kindly provided by C. Bargmann, Rockefeller University, New York, USA). For *nmur-1*, the *nmur-1* cDNA was first amplified from mixed-stage wild-type *C. elegans* template and inserted into a pSM backbone (linearized by BamHI and NheI restriction digest) using Gibson assembly. The 1kb *nmur-1* 3’UTR sequence was then inserted after the GFP coding sequence by Gibson assembly into the backbone digested by EcoRI and SpeI, which replaced the *unc-54* 3’UTR sequence native to the original pSM plasmid. Finally, a 2 kb *nmur-1* promoter fragment was inserted prior to the gene’s cDNA sequence after SphI and XbaI digestion.

For cell-specific rescue experiments, the *nmur-1::SL2::gfp::nmur-1 3’UTR* pSM backbone was first linearized by PCR amplification, yielding a 5900 bp long linear sequence in which promoters driving expression in selected neurons were inserted using Gibson Assembly. For this, promoter regions were amplified from genomic N2 DNA, with a sequence 2877 bp upstream of the predicted coding region for *gcy-28d*; 2648 bp for *odr-2*; 2393 bp for *osm-6*; 3000 bp for *ceh-23*; 2534 bp for *ocr-2*; 2009 bp for *gcy-8*; 2067 bp for *srh-142*; 2449 bp for *srh-220* and 4786 bp for *ocr-4*. In addition, a 314 bp *ins-1(s)p* was amplified from pCEC19 (Bargmann lab) and a 485 bp *flp-1(tr)p* from pCS178 (Gottschalk lab).

For *capa-1* expression analysis, first the 2.5 kb long *capa-1* 3’UTR sequence was inserted into the pSM backbone linearized by EcoRI and SpeI using Gibson assembly. Next, a genomic region encompassing 2.2 kb of the sequence upstream of the *capa-1* start codon and the *capa-1* gene sequence were amplified from genomic DNA of mixed-stage wild-type *C. elegans* and inserted into the pSM backbone by Gibson assembly (backbone digested by BamHI and NheI).

For ASG-specific silencing using Arch or tetanus toxin (TeTx), we inserted the 2.2 kb *capa-1* promoter sequence upstream of the Arch and TeTx coding sequences in pSH122 (linearized by SphI and MscI) and pSH188 (linearized by NotI and NheI), respectively.

The transgenes for ASG-specific *capa-1* knockdown were constructed as previously described^4^. Briefly, the *ops-1* promoter (1.9 kb) was fused to a 1132 bp genomic *capa-1* fragment in sense and antisense orientation. The genomic *capa-1* fragment is identical to that of the clone taken up in the Ahringer feeding RNAi library^5^.

The transgene for ASG-specific *gcy-15p::mCherry expression* was constructed through fusion PCR. First, the 995 bp mCherry sequence was amplified from pGH8, the 1908 bp promoter and the 883 bp 3’UTR of the *gcy-15* gene amplified. All three fragments were sequentially fused through fusion PCR.

### *In vitro* GPCR activation assay

The GPCR activation assay was performed as previously described^6,7^. Briefly, mammalian CHO K1 cells stably overexpressing apo-aequorin and human Gα_16_ were transiently transfected with the *nmur-1*/pcDNA3.1 or empty pcDNA3.1 (control) plasmid using Lipofectamine LTX and Plus reagent (Invitrogen). After transfection, cells were grown overnight at 37 °C, after which they were transferred to 28 °C and allowed to incubate for 24 h. On the day of the assay, CHO cells were harvested from their culture flask and loaded with Coelenterazine H (Invitrogen) for 4 h at room temperature, which elicits the formation of the calcium-sensitive photoprotein aequorin. The incubated cells were then added to synthetic peptides dissolved in DMEM/BSA, and luminescence measured for 30 s at 496 nm using a Mithras LB940 luminometer (Berthold Technologies). After 30 s of readout, 0.1 % triton X-100 was added to lyse the cells, resulting in a maximal calcium response that was measured for 10 s. After initial screening, concentration-response curves were constructed for HPLC-purified CAPA-1 peptides by subjecting the transfected cells to each peptide in a concentration range from 0.1 nM to 100 µM. Cells transfected with an empty vector were used as a negative control. Assays were performed in triplicate on at least two independent days. Concentration-response curves were fitted using GraphPad Prism 5 (nonlinear regression analysis with a sigmoidal concentration-response equation).

A peptide library of over 300 synthetic *C. elegans* peptides that was used to challenge the NMUR-1 expressing CHO cells was compiled based on *in silico* predictions and peptidomics data^8,9^. Peptides were synthesized by Thermo Scientific and GL Biochem Ltd.

### Transgenesis and expression pattern analysis

To generate transgenic *C. elegans*, constructs were injected into the syncytial gonad of young adult worms at concentrations ranging from 5 to 50 ng/µL with 50 ng/µL of the coinjection marker *elt-2p::GFP* or *unc-122p::dsRED* and 17 ng/µL of a 1-kb DNA ladder (Thermo Scientific) as carrier DNA.

Expression patterns of reporter transgenes were visualized by an inverted Zeiss AxioObserver Z1 microscope fitted with a 40X oil objective, an ORCA-Flash4.0 V2 camera (Hamamatsu) and W-View GEMINI image splitting optics (Hamamatsu). Image acquisition was performed using Metamorph (Molecular Devices) software. For imaging, hermaphrodite animals were immobilized on 10% agarose pads using polystyrene beads between pad and coverslip to restrain the animal’s movement (Polybead® Microspheres 0.10μm, Polyscience). Expression patterns were confirmed in at least two independent transgenic strains.

### Salt chemotaxis assays

Salt chemotaxis and gustatory plasticity behavior were assessed as described previously^10,11^ (Figure 1). All behavioral assays were performed in a climate-controlled room set at 20°C and 40% relative humidity. In brief, 1-day young adult hermaphrodites were grown at 25°C on culture plates seeded with sufficient *E. coli* OP50 or HT115. For chemotaxis towards salt, worms were collected from their culture plate and washed three times over a period of 15 minutes with chemotaxis buffer (CTX, 5 mM KH_2_PO_4_/K_2_HPO_4_ pH 6.6, 1 mM MgSO_4_, and 1 mM CaCl_2_). The attraction to salt is then assessed by pipetting 100 – 200 animals on the intersection of four-quadrant plates (Falcon X plate, Becton Dickinson Labware) filled with buffered agar (2% agar, 5 mM KH_2_PO_4_/K_2_HPO_4_ pH 6.6, 1 mM MgSO_4_, and 1 mM CaCl_2_) of which two opposing pairs have been supplemented with NaCl (100, 200, 300 or 400 mM NaCl). Assay plates were always prepared fresh and left open to solidify and dry for 60 min. Plates were then closed and used on the same day. After animals were allowed to crawl for 10 minutes on the quadrant plate, a chemotaxis index was calculated as (n(A) – n(C)) / (n(A) + n(C)) where n(A) is the number of worms within the quadrants containing NaCl and n(C) is the number of worms within the control quadrants without NaCl.

For gustatory plasticity, the well-fed synchronized adult worm population is washed in CTX buffer with or without 100 mM NaCl for NaCl-conditioned and mock-conditioned worms, respectively. The response to 25 mM NaCl is then assayed on four-quadrant plates to which 25 mM NaCl has been added to one pair of opposing quadrants. After 10 min, the distribution of worms over the quadrants is determined and a chemotaxis index calculated as described above.

For tracking analysis, conditioned behavior is assessed on 0 to 100 mM linear NaCl gradients^12^. These are generated in square Petri dishes (Greiner, 120 x 120 x 17mm, vented), by elevating one side of the plate and filling half of the plate with 50 mL of buffered agar supplemented with 100 mM NaCl. Allowing the agar to solidify for 30 min creates a triangle wedge, after which the plate is laid flat and the other half of the plate is filled with 50 mL buffered agar without NaCl. Behavioral assays were performed 24 hr after the gradients were established. In each assay, 15–20 young adult worms were mock- or NaCl-conditioned as described above, and pipetted to the middle of the square plate corresponding to approximately 50 mM NaCl.

For pre-exposure to NaCl in the presence of food^13^, adult animals were washed off culture plates using CTX buffer with or without 100 mM NaCl and allowed to sediment for a few minutes. They are then transferred to agar plates filled with buffered agar (2% agar, 5 mM KH_2_PO_4_/K_2_HPO_4_ pH 6.6, 1 mM MgSO_4_, and 1 mM CaCl_2_), with or without 100 mM NaCl, unseeded or seeded with 200 µL of 0.5 g/mL freshly grown OP50 bacteria in H_2_O. After 30 minutes of pre-exposure, animals were washed of the plates using CTX buffer ± NaCl, allowed to sediment for 1 min, and placed on a chemotaxis quadrant plate (25 mM NaCl) to measure salt chemotaxis behavior.

### Behavioral Quantification

Assay plates were imaged using an in-house tracking platform consisting of 10 MP NET GP11004M cameras fitted with LM16JC10M KOWA lenses. A diffuse LED light source (Rosco 12”x12” LitePad) is used to illuminate animals in trans, while two consumer privacy filters (3M PF17.0) are stacked perpendicularly on the glass stage holding the assay plates to enhance contrast. StreamPix 6 multicamera software was used to acquire footage at 2 frames per second. Because behavioral assays are sensitive to ambient conditions, care was taken to record experimental and control conditions simultaneously by dedicated cameras.

The resulting footage was analyzed using custom particle-tracking algorithms written in MATLAB (Mathworks, Natick, MA), built upon the algorithm of a previously reported script^14^. Briefly, worms are identified by average background subtraction, gaussian smoothing of the resulting foreground image and intensity thresholding to binarize each frame. Blobs falling within a predefined size range are connected over adjacent frames into individual tracks, and coordinates of the center of mass (‘centroid’) and simple shape parameters are saved for each worm object on each frame. Individual worm tracks are terminated when worms collide with each other or with the boundaries of the behavioral arena, while tracks with a total duration of less than 5 min are discarded. The resulting trajectories were smoothed with a 1.5 second moving average filter before calculating instantaneous speed of the centroid.

The movement of each animal’s center of mass was considered to automatically label different behavioral states. Pauses are labelled when the centroid’s velocity falls under 0.01 mm/s for more than half of all time points in a 10 sec sliding time window. The animal’s behavioral state is classified as a turn if, for any given time point, the geometric angle between the points behind or ahead the track by 0.3 mm is less than 80°^15^. Any time points in which the velocity remains smaller than 0.1 mm/s after the turn are also classified as turn^16^. According to this definition, reversals and omega turns are both recognized as turns^17^. When individual turns are separated by less than 3.8 sec, they are bundled into a pirouette^15^.

To quantify NaCl attraction of mock- or NaCl-conditioned worms on linear NaCl gradients, several indices were computed. For each frame in the 30-minute recording after release on the linear NaCl gradient, the position of each individual worm is averaged, and the population mean position traced through time with shaded regions representing S.E.M. For statistical comparison, the average position of each worm within the defined time interval is computed per condition. Chemotactic rate and direction is quantified for each individual animal trajectory by dividing the mean velocity along the gradient direction 〈v_g_〉 by the mean crawling speed along the trajectory 〈s〉. This index is positive when an individual worm predominantly moves up the gradient, and negative when it migrates to lower salt concentrations. To quantify the biased random walk index, trajectory stretches in which worms exhibiting run behavior are extracted, and the fractional difference of run duration up or down the gradient is calculated for each worm (〈*rup*〉−〈*rdown*〉)/(〈*rup*〉+〈*rdown*〉). The klinotaxis index is determined by the fractional difference in the probability of run behavior after a reorientation event (turn of pirouette) to be initiated up or down the gradient (〈*pup*〉−〈*pdown*〉)/(〈*pup*〉+〈*pdown*〉) ^12^.

### Locomotion on a bacterial lawn and after removal from food

For quantitative analyses of local search behavior and roaming or dwelling, worms were grown until the first day of adulthood at a low population density, on NGM plates seeded with OP50 or HT115, in a climate-controlled room (20°C and 40% relative humidity). For assays on food, assay plates were prepared one day prior to the experiment by seeding freshly poured 90 mm diameter Petri plates with 500 µL of fresh OP50 or HT115 culture and allowing the bacteria to grow overnight in the climate-controlled room. On the day of the assay, 5 to 10 animals were first picked to a holding plate prepared in parallel with the assay plates. After 45 min, animals were moved to the assay plate and placed on the imaging platform and recorded for 90 min after 30 min of acclimatization. Worms were tracked from this footage as described in the previous section, discarding tracks shorter than 20 min. To classify behavior into roaming and dwelling, speed and angular speed (a measure of path curvature) were averaged over 10 s intervals. Both classes are separated by tracing a line (Speed = Curvature/450) through the plot scattering speed in function of curvature. Time points falling above the line were considered roaming intervals, while those below the line were labelled as dwelling^18,19^. For each animal, the fraction of time in each behavioral state was computed, as well as each individual’s average speed during dwelling or roaming. For local search behavior off food, 5 to 10 well-fed worms were picked from their culturing plate to an unseeded intermediate plate and within 1 min transferred to another unseeded plate on which they were filmed for 15 min ^17^. Mean speed was calculated per individual, as well as the fraction of time spent in each behavior as determined by the segmentation algorithm explained above.

### Imaging of neuronal activity in microfluidic chip

Transgenic worms expressing GCaMP3 in ASG^20^ or YC2.12 in ASE neurons^21^ were imaged in a microfluidic polydimethylsiloxane (PDMS) chip^22^. This PDMS chip is attached to inlet tubes, connecting the worm-loading channel to a reservoir of CTX buffer (with or without 100 mM NaCl). Well-fed transgenic worms were first picked in the reservoir, after which they were exposed to the CTX buffer in the absence of food for approximately 15 minutes. NaCl-responses were then assayed by positioning the worm in the channel and challenging the worm to either 0 to 50 mM NaCl upshift or 50 to 0 mM NaCl downshift. The composition of the buffers for imaging is the same as the buffers used in the behavioral assay, except for the addition of sucrose to balance osmolarity (5 mM KH_2_PO_4_/K_2_HPO_4_ pH 6.6, 1 mM MgSO_4_, and 1 mM CaCl_2_ and 0, 50 or 100 mM NaCl and adjusted to 350 mOsmol/kgH_2_O by sucrose). Imaging was conducted on an inverted Zeiss AxioObserver Z1 microscope. Fluorescent images were captured using a 40X objective and an ORCA-Flash4.0 V2 camera (Hamamatsu) driven by Metamorph software (Molecular Devices). Images were analyzed using custom-written Mathematica (Wolfram) code setting a region of interest around the cell body. For GCaMP3 imaging, the adjacent background was subtracted on each image and the fluorescence intensity during the 10 second period before stimulus delivery was averaged and defined as F_0_. ΔF/F0 (%) is calculated as 100*(background corrected fluorescence – F_0_) / F_0_. For YC2.12 imaging, ΔR/R (%) is calculated as the ratio between CFP and YFP fluorescent emissions.

### Optogenetic silencing of ASG

Transgenic worms expressing the outward-directed proton pump Arch in ASG were cultured at 25°C on culture plates seeded with sufficient OP50 *E. coli*. One day before the assay, transgenic worms were transferred to OP50 culture plates to which *all-trans* retinal (ATR, Sigma Aldrich) has been added (300 µM final concentration in fresh OP50 liquid culture). This cofactor is needed for Arch activity, and allows transgenic worms transferred to culture plates without ATR to serve as negative control. A 0 to 50 mM linear gradient was generated in a 55 mm Petri plate conform to the protocol for linear gradients in square plates (15 mL buffered agar supplemented with 50 mM NaCl is poured in a 55 mm diameter Petri dish elevated to one side. An additional 15.5 mL buffered agar without NaCl is poured on top of the first triangular wedge once it is hardened. NaCl is allowed to diffuse at least 24 h prior the assay). Individual worms were washed for 15 min in a small volume of CTX buffer with or without 100 mM NaCl (NaCl-conditioning or mock-conditioning). After conditioning, the worm was transferred to the middle of the 0 to 50 mM NaCl gradient plate mounted on a Zeiss AxioObserver Z1 microscope. The worm was then allowed to navigate the plate for 10 min while an automated xy-stage kept it under in the field of view. For this, a low-resolution Stingray F-145B camera (Allied Vision) was mounted to the microscope, and repositioned the Prior ProScan III stage each second to keep the worm in the middle of the field of view using Matlab code adapted from Wang et al., 2013^23^. Each animal’s path was reconstructed based on the recorded low-resolution video-stream and the logged stage positions. The behavior was then segmented and indices reflecting chemotaxis behavior were calculated as described above. For ASG silencing during conditioning, the worm was kept under the yellow-green light under a fluorescent stereomicroscope during the washing procedure (Leica MZ16F microscope and EL6000 external light source with mCherry Leica filter set, number 10450195 – 540 - 580 nm. Intensity approximately 0.2 to 0.4 mW/mm²). For silencing during the navigation period, the worm was continuously illuminated with 567 – 602 nm light while it was being tracked (Zeiss Colibri.2 LED module with Filter cube 61 HE GFP/HcRed, intensity approximately 0.552 mW/mm²).

### Quantification and statistical analysis

Behavioral tracking and analysis of chemotaxis indices was performed in MATLAB (MathWorks). For salt chemotaxis assays, statistical significance of the mean position of the population on the gradient and the behavioral indices was determined using one-way ANOVA and Tukey post-hoc for multiple comparisons. The time interval for which these indices are calculated is indicated with a horizontal black bar on the corresponding mean population position through time plot. Shaded bars around the mean population traces denote S.E.M. For boxplots on which individual data points are scattered, the central mark indicates the median and the bottom and top edges the 25th and 75th percentiles, respectively. The whiskers extend to the most extreme data points not considered as outlier. For the fraction of time in each behavioral state, the relative fraction was binned over individual worms and analyzed for statistical significance by Chi-squared analysis. The time spent in roaming or dwelling states while foraging on a bacterial lawn was analyzed using a Kruskal-Wallis test. For chemotaxis studies in which worms were pre-exposed to NaCl in the presence or absence of food, a two-way ANOVA was used in combination with a Tukey post-hoc test. Data from optogenetic experiments was analyzed using a three-way ANOVA followed by a Tukey post-hoc test.

All behavioral assays were conducted at least four times on at least two separate days. For transgenic experiments, animals with clear target fluorescence emanating from the pSM backbone, TeTx::mCherry or Arch::YFP fusions were selected as transgenic animals. Siblings without this fluorescence (or not having fluorescence from the co-injection marker in the case of *nmur-1* rescue and ASG-specific *capa-1* RNAi experiments) were used as control and showed behavior similar to non-transgenic mutant animals.

## Supplementary figure legends

**Supplementary figure 1.**
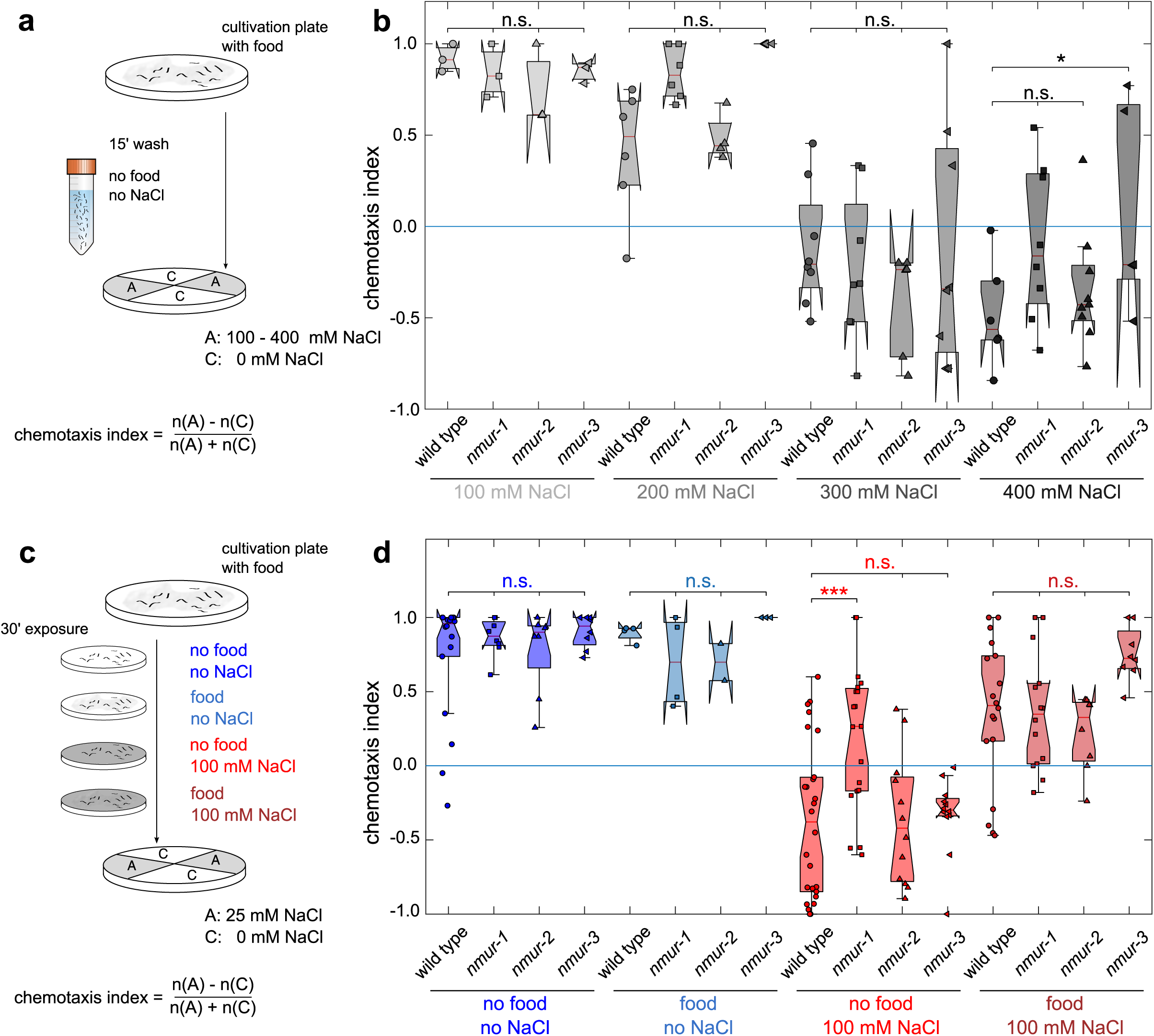
Mutants of *nmur-1* display normal salt sensing but are defective in associating salt with the lack of food (related to Figure 1) **(a)** Overview of chemotaxis assay for behavioral responses to increasing NaCl concentrations. Synchronized adult *C. elegans* are washed from culture plates and washed for 15 minutes in NaCl-free buffer. After washing, the attraction to salt is assayed on quadrant plates of which two opposing quadrants are supplemented with 100, 200, 300 or 400 mM NaCl. The distribution of animals on the quadrants is determined at 10 minutes and a chemotaxis index calculated. **(b)** Mean chemotaxis index of wild type animals and *nmur-1*, *nmur-2* and *nmur-3* mutants to increasing NaCl concentrations. n ≥ 3 assays per genotype. **(c)** Overview of NaCl chemotaxis assay under different conditioning regimes. Synchronized adult *C. elegans* are washed from culture plates and conditioned for 30 min on agar plates supplemented with 100 mM NaCl. Conditioning plates without NaCl and/or seeded with *E. coli* OP50 bacteria are used as a control. After conditioning, the attraction to NaCl is assayed on quadrant agar plates in which two opposing quadrants contain 25 mM NaCl. A chemotaxis index is calculated as the fractional proportion of worms attracted to salt. **(d)** Mock-conditioned animals, which have not been pre-exposed to salt (with or without food) show strong attraction to NaCl. Conditioning with salt in the absence of food switches salt chemotaxis behavior to an avoidance response, but this conditioned response is significantly reduced in *nmur-1* mutant animals. The *nmur-1* mutant only shows defects in gustatory learning when worms are pre-exposed to NaCl in the absence of food. *nmur-1* animals have normal salt chemotaxis behavior when they are conditioned with salt in the presence of food, which indicates that they are defective in gustatory aversive learning. n ≥ 3 assays per genotype.

**Supplementary figure 2.**
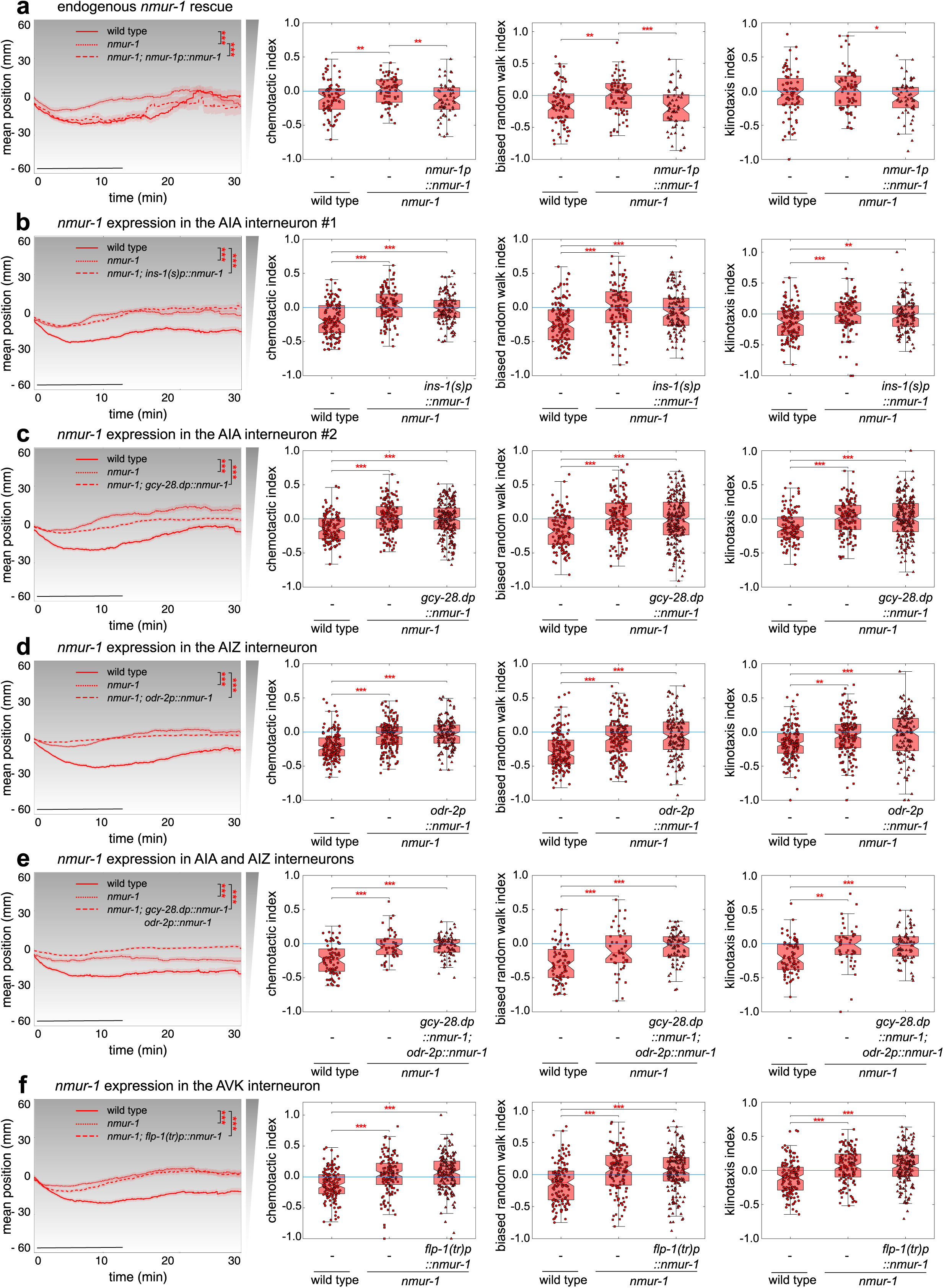
Endogenous and cell-selective *nmur-1* rescue in selected interneurons (related to Figure 2) Mean position on the NaCl-gradient, and chemotactic, biased random walk and klinotaxis indices for NaCl-conditioned *nmur-1* mutant worms in which *nmur-1* expression is specifically restored **(a)** under the control of its promoter (*nmur-1p::nmur-1* genomic sequence). n ≥ 50 animals per genotype. **(b)** in the AIA interneuron under control of the *ins-1(s)* promoter sequence (*ins-1(s)p::nmur-1* cDNA^1^). n ≥ 161 animals per genotype. **(c)** in the AIA interneuron under the control of the *gcy-28d* promoter sequence (*gcy-28dp::nmur-1* cDNA^2^). n ≥ 119 animals per genotype. **(d)** in the AIZ interneuron under the control of the *odr-2* promoter sequence (*odr-2p::nmur-1* cDNA^3^). n ≥ 146 animals per genotype. **(e)** in AIA and AIZ interneurons under the control of the *gcy-28d* and *odr-2* promoter sequence, respectively (*gcy-28dp::nmur-1* cDNA; *odr-2p::nmur-1* cDNA) n ≥ 47 animals per genotype. **(f)** in the AVK interneuron under the control of the *flp-1(tr)* promoter sequence (*flp-1(tr)p::nmur-1* cDNA^4^). n ≥ 136 animals per genotype. Black bar on mean position plot (left panel) indicates the time interval used for statistical comparison of the mean position on the gradient, but also for calculating the chemotactic, biased random walk and klinotaxis indices. Shaded area around mean position traces denotes S.E.M. Individual datapoints are scattered on boxplots (red line marks the median, boxes indicate the 25^th^ to 75^th^ percentile, whiskers extend to the extreme datapoints not considered an outlier). Statistical comparisons by two-way ANOVA and Tukey post-hoc test. * P<0.05; ** P<0.01; *** P<0.001.

**Supplementary figure 3.**
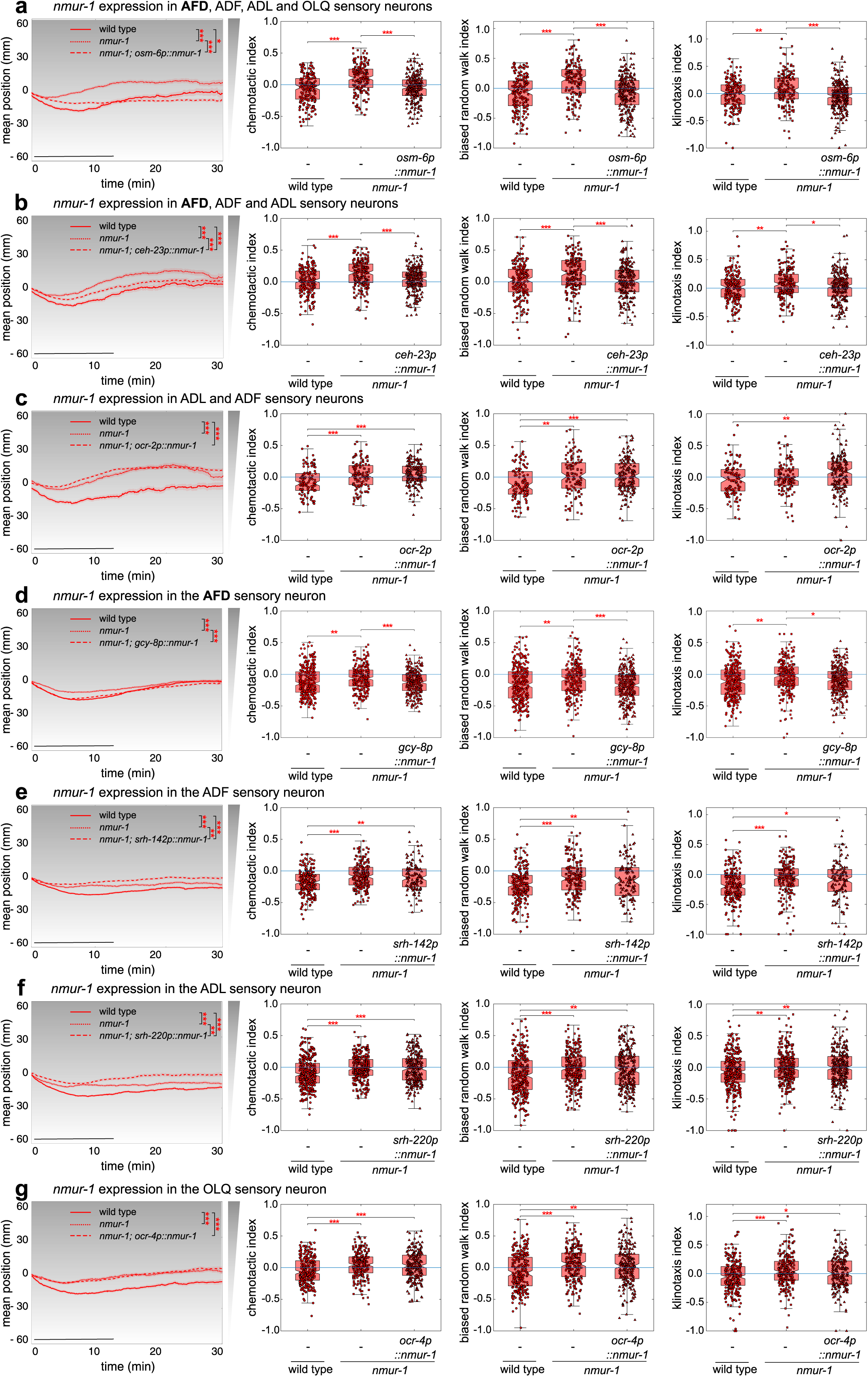
Cell-selective *nmur-1* rescue in selected sensory neurons (related to Figure 2) Mean position on the NaCl-gradient, and chemotactic, biased random walk and klinotaxis indices for NaCl-conditioned *nmur-1* mutant worms in which *nmur-1* expression is specifically restored **(a)** in AFD, ADF, ADL and OLQ sensory neurons under the control of the *osm-6* promoter sequence (*osm-6p::nmur-1* cDNA^5^). n ≥ 170 animals per genotype. **(b)** in AFD, ADF and ADL sensory neurons under the control of the *ceh-23* promoter sequence (*ceh-23p::nmur-1* cDNA^6^). n ≥ 185 animals per genotype. **(c)** in ADF and ADL sensory neurons under the control of the *ocr-2* promoter sequence (*ocr-2p::nmur-1* cDNA^7^). n ≥ 152 animals per genotype. **(d)** in the AFD sensory neuron under the control of the *gcy-8* promoter sequence (*gcy-8p::nmur-1* cDNA^8^). n ≥ 284 animals per genotype. **(e)** in the ADF sensory neuron under the control of the *srh-142* promoter sequence (*srh-142p::nmur-1* cDNA^9^). n ≥ 127 animals per genotype. **(f)** in the ADL sensory neuron under the control of the *srh-220* promoter sequence (*srh-220p::nmur-1* cDNA^10^). n ≥ 275 animals per genotype. **(g)** in the OLQ sensory neuron under the control of the *ocr-4* promoter sequence (*ocr-4p::nmur-1* cDNA^7^). n ≥ 223 animals per genotype.

**Supplementary figure 4.**
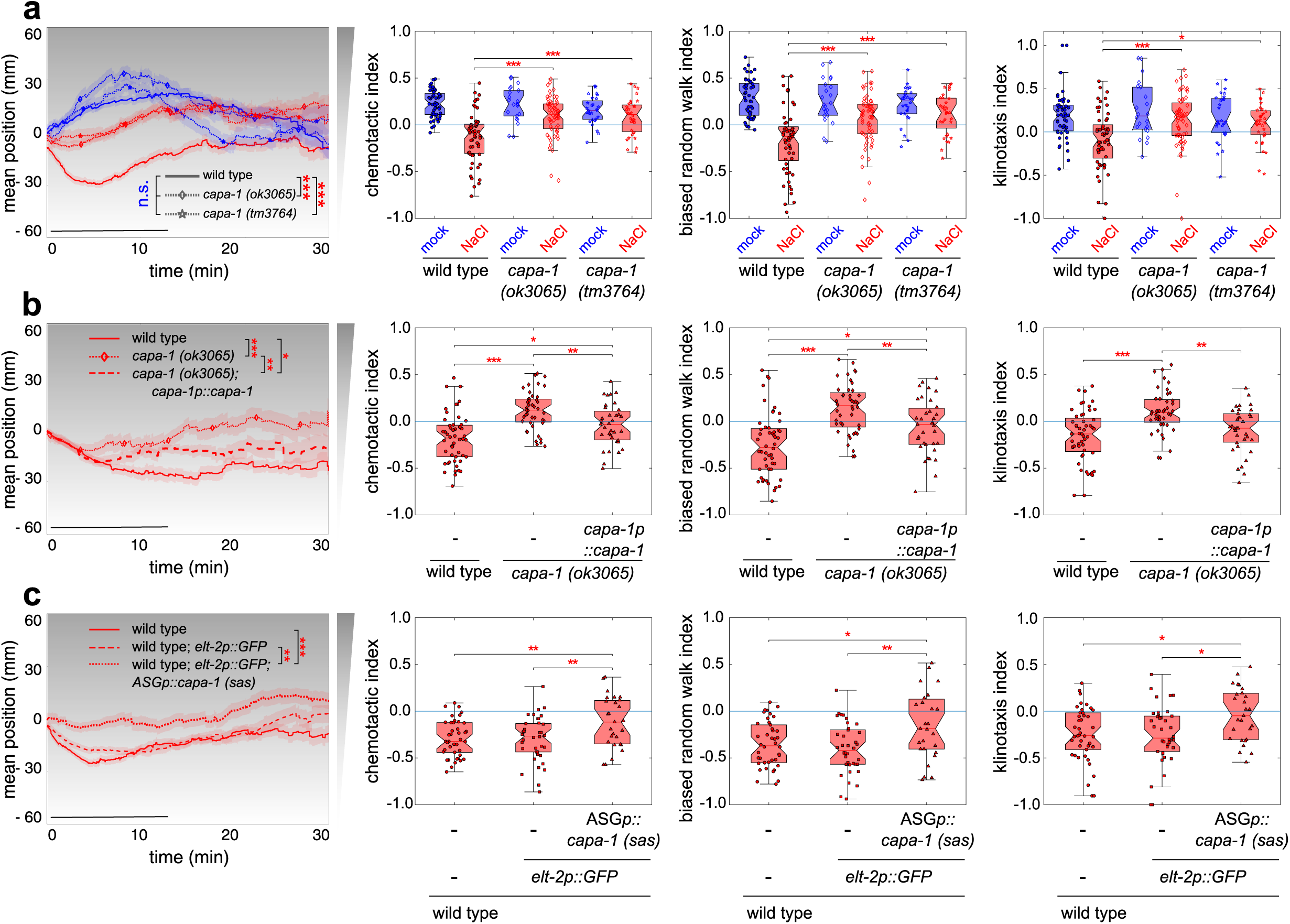
CAPA-1 signaling from ASG neurons regulates the experience-dependent modulation of NaCl chemotaxis behavior (related to Figure 3) Mean position on the NaCl-gradient, and chemotactic, biased random walk and klinotaxis indices for **(a)** two independent deletion mutants of *capa-1* after mock- and NaCl-conditioning. n ≥ 25 animals per genotype. **(b)** NaCl-conditioned *capa-1 (ok3605)* mutant worms in which *capa-1* expression is restored under the control of its endogenous promoter sequence (*capa-1p::capa-1*). n ≥ 37 animals per genotype. **(c)** NaCl-conditioned wild-type worms in which *capa-1* is specifically silenced in ASG (*ops-1p::capa-1 sense and antisense*^11^). n ≥ 27 animals per genotype.

**Supplementary figure 5.**
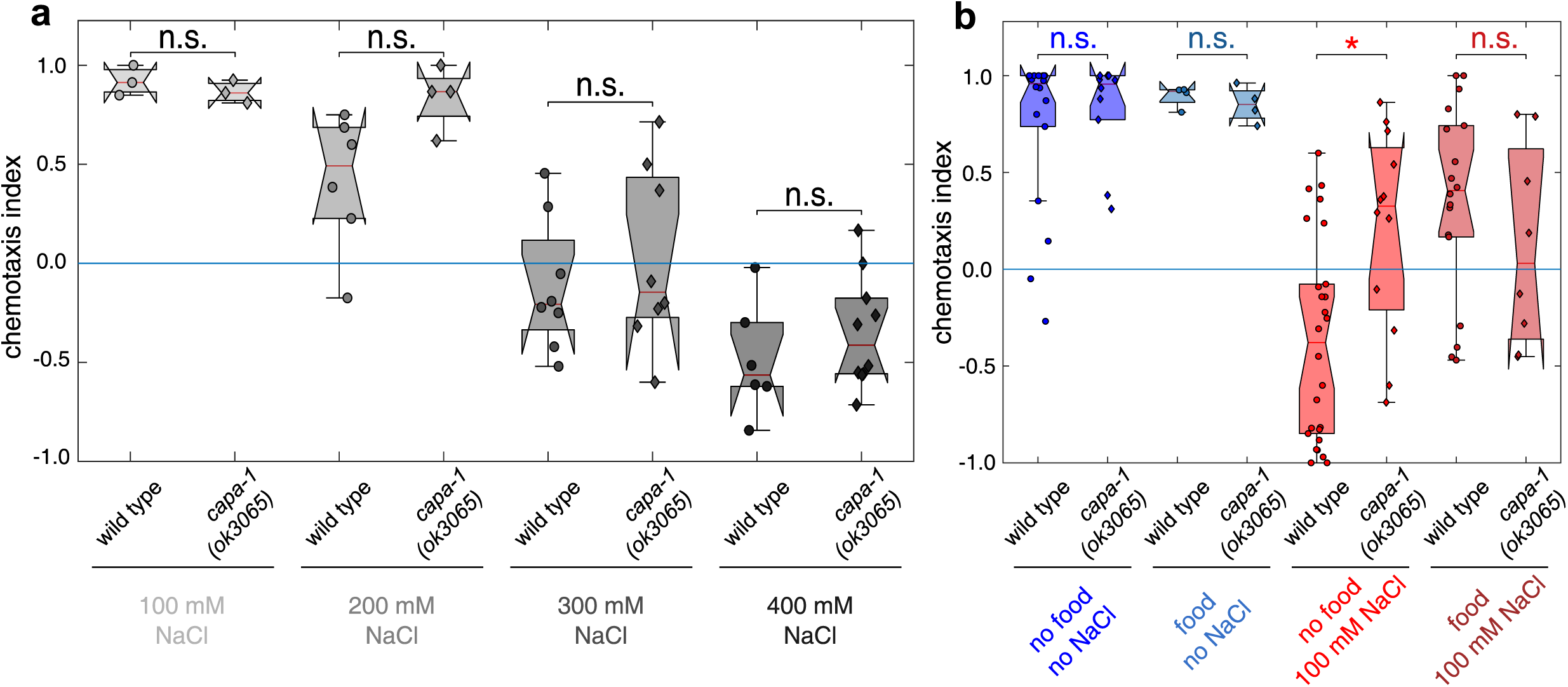
*capa-1* Mutants Have Wild-type Sensory Responses to NaCl, But Are Defective in Gustatory Associative Learning (related to Figure 3) **(a)** Mean chemotaxis index (according to Supplementary figure 1a) of wild type and *capa-1 (ok3065)* mutant animals to increasing NaCl concentrations. n ≥ 4 assays per genotype. **(b)** NaCl chemotaxis behavior of wild type and *capa-1 (ok3065)* mutant animals under different conditioning regimes (according to Supplementary figure 1c). n ≥ 3 assays per genotype.

**Supplementary figure 6.**
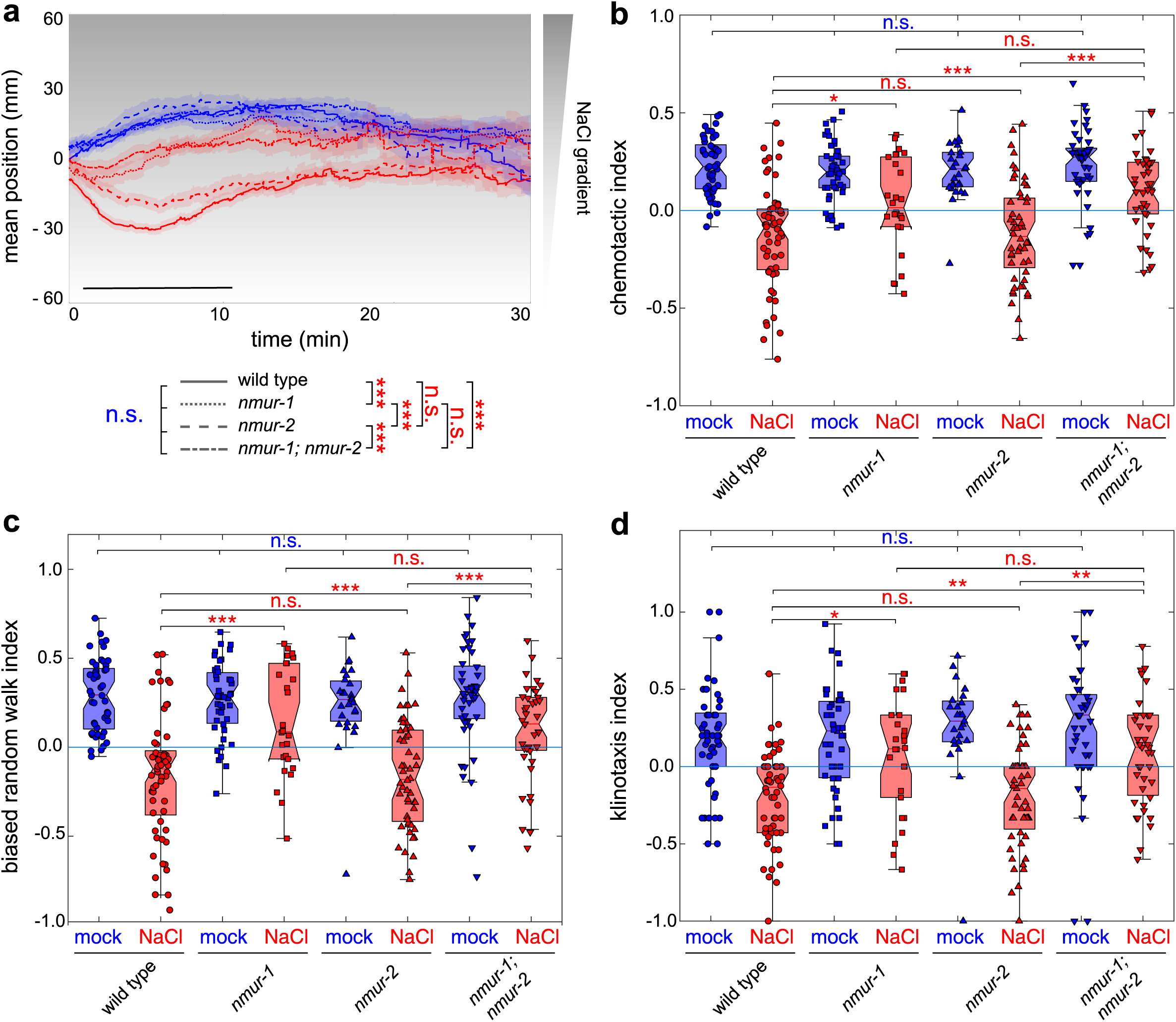
The CAPA-1 receptor NMUR-2 is not involved in gustatory associative learning. **(a)** Mean position of *nmur-1* and *nmur-2* single and double mutants on a 0 - 100 mM NaCl gradient after mock- (blue traces) or NaCl-conditioning (red traces). Shaded regions represent S.E.M. Black bar indicates the time interval used for statistical comparison of the mean position on the gradient, but also for calculating the chemotactic, biased random walk and klinotaxis indices in panels b-d. **(b)** Chemotactic, **(c)** biased random walk and **(d)** klinotaxis indices for *nmur-1* and *nmur-2* single and double mutants after mock- and NaCl-conditioning. n ≥ 27 animals per genotype.

**Supplementary figure 7.**
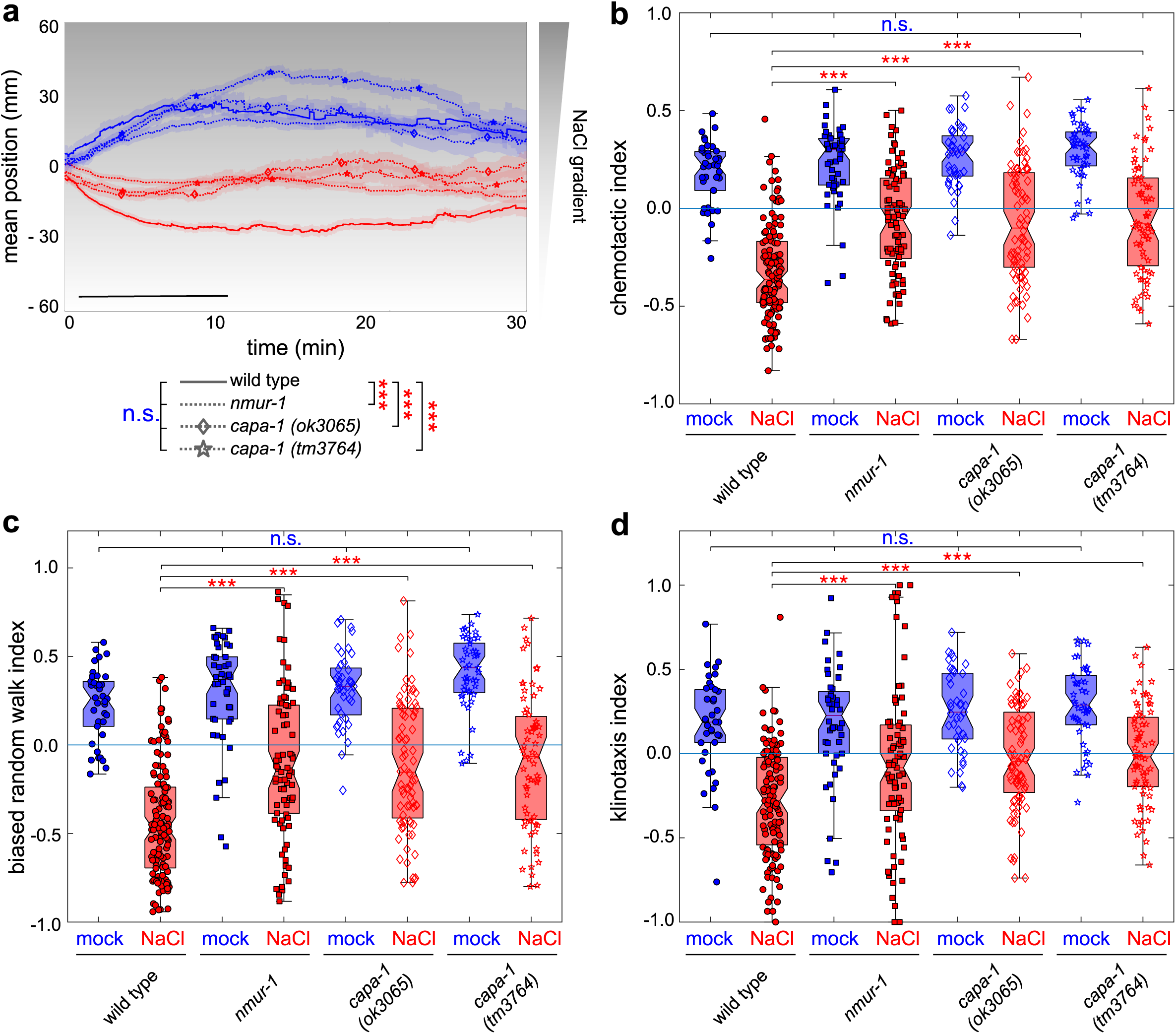
Defects in gustatory associative learning of *capa-1* and *nmur-1* mutants are independent of the bacterial diet. **(a)** Mean position on a NaCl gradient for mock- and NaCl-conditioned wild type, *capa-1* and *nmur-1* animals cultivated on *E. coli* HT115 bacteria. Shaded regions around the mean traces denote S.E.M. Black bar indicates the time interval used for statistical comparison of the mean position on the gradient, but also for calculating the chemotactic, biased random walk and klinotaxis indices in panels b-d. **(b)** Chemotactic, **(c)** biased random walk and **(d)** klinotaxis indices for wild type animals, *capa-1* and *nmur-1* mutants cultivated on *E. coli* HT115. n ≥ 49 animals per genotype.

**Supplementary figure 8.**
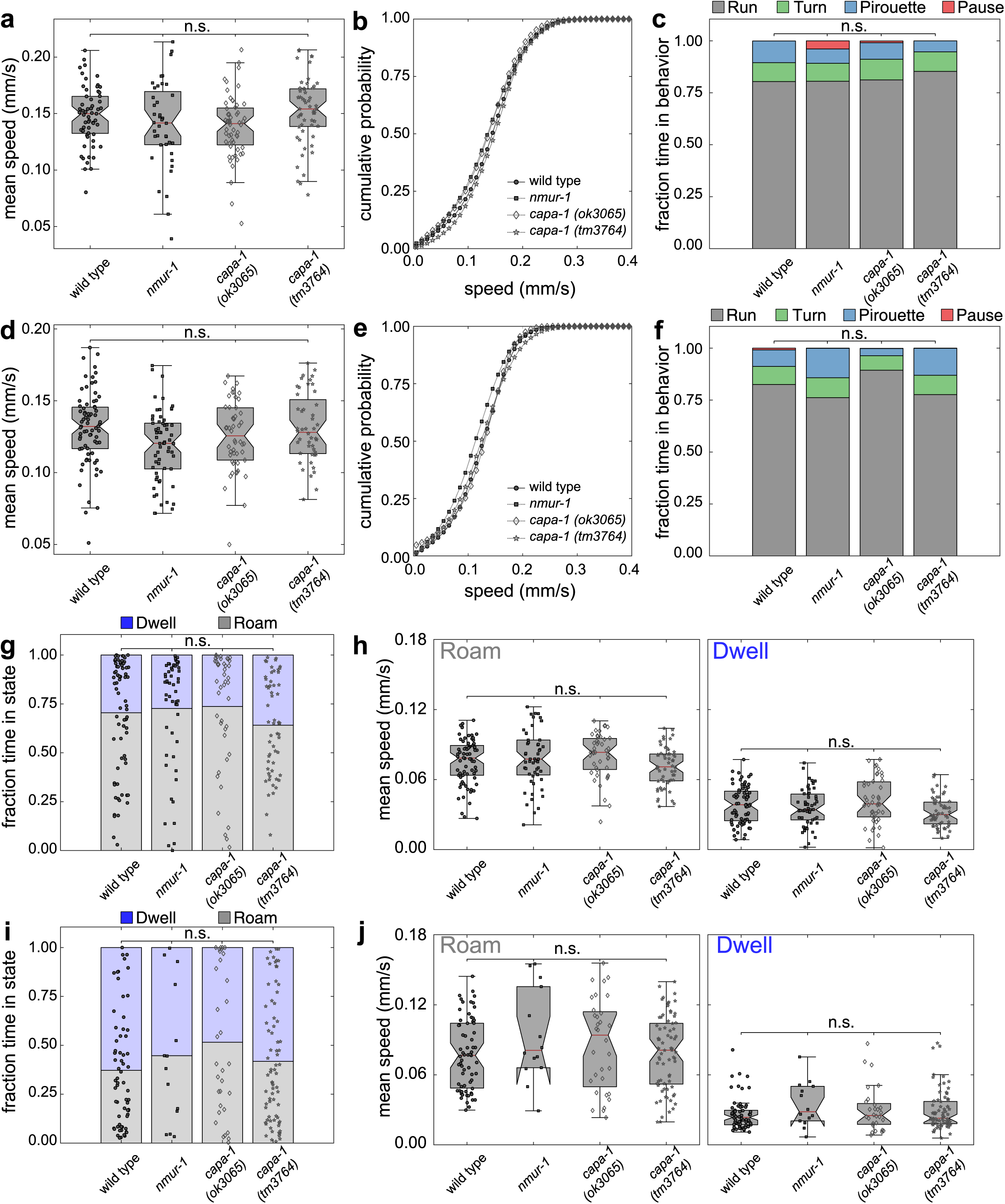
Mutants defective in CAPA-1 signaling show normal food searching behaviors. **(a - c)** Local search behavior of worms grown on *E. coli* OP50. Adult wild type, *nmur-1* and *capa-1 C. elegans* are removed from OP50 and transferred to an unseeded agar plate for behavioral tracking. **(a)** Average speed and **(b)** cumulative probability distribution of speed after removal from food. **(c)** Individual worm trajectories from are subdivided into discrete stretches of Run (gray) – Turn (green) – Pirouette (blue) or Pause (red) behavior, and average fraction of time in each behavioral state plotted. n ≥ 40 animals per genotype. **(d - f)** Local search behavior of worms grown on *E. coli* HT115. Adult wild type, *nmur-1* and *capa-1 C. elegans* are removed from OP50 and transferred to an unseeded agar plate for behavioral tracking. **(d)** Average speed and **(e)** cumulative probability distribution of speed after removal from food. **(f)** Individual worm trajectories from are subdivided into discrete stretches of Run (gray) – Turn (green) – Pirouette (blue) or Pause (red) behavior, and average fraction of time in each behavioral state plotted. n ≥ 51 animals per genotype. **(g and h)** Quantification of roaming and dwelling behaviors on *E. coli* OP50. While feeding on a bacterial lawn, *C. elegans* alternates between two distinct behavioral states: an active exploratory state referred to as roaming, and more passive behavior termed dwelling. Both states are assigned based on locomotion speed and angular speed (a measure of turning rate). Roaming animals move quickly across the bacterial lawn and turn infrequently to explore the bacterial lawn, while dwelling animals remain in a small area by moving slowly and turning more frequently. **(g)** The fraction of time spent in the roaming state and **(h)** average speed during roaming or dwelling for wild type animals and *capa-1* or *nmur-1* mutants on OP50. n ≥ 44 animals per genotype. **(i and j)** Quantification of roaming and dwelling behaviors on *E. coli* HT115. **(i)** The fraction of time spent in the roaming state and **(j)** average speed during roaming or dwelling is similar for wild type animals and *capa-1* or *nmur-1* mutants. n ≥ 14 animals per genotype. Statistical comparisons by one-way ANOVA and Tukey post-hoc test (a, d, h and j), Chi-square test on binned data (c and f), and Kruskal-Wallis test (g and i).

**Supplementary figure 9.**
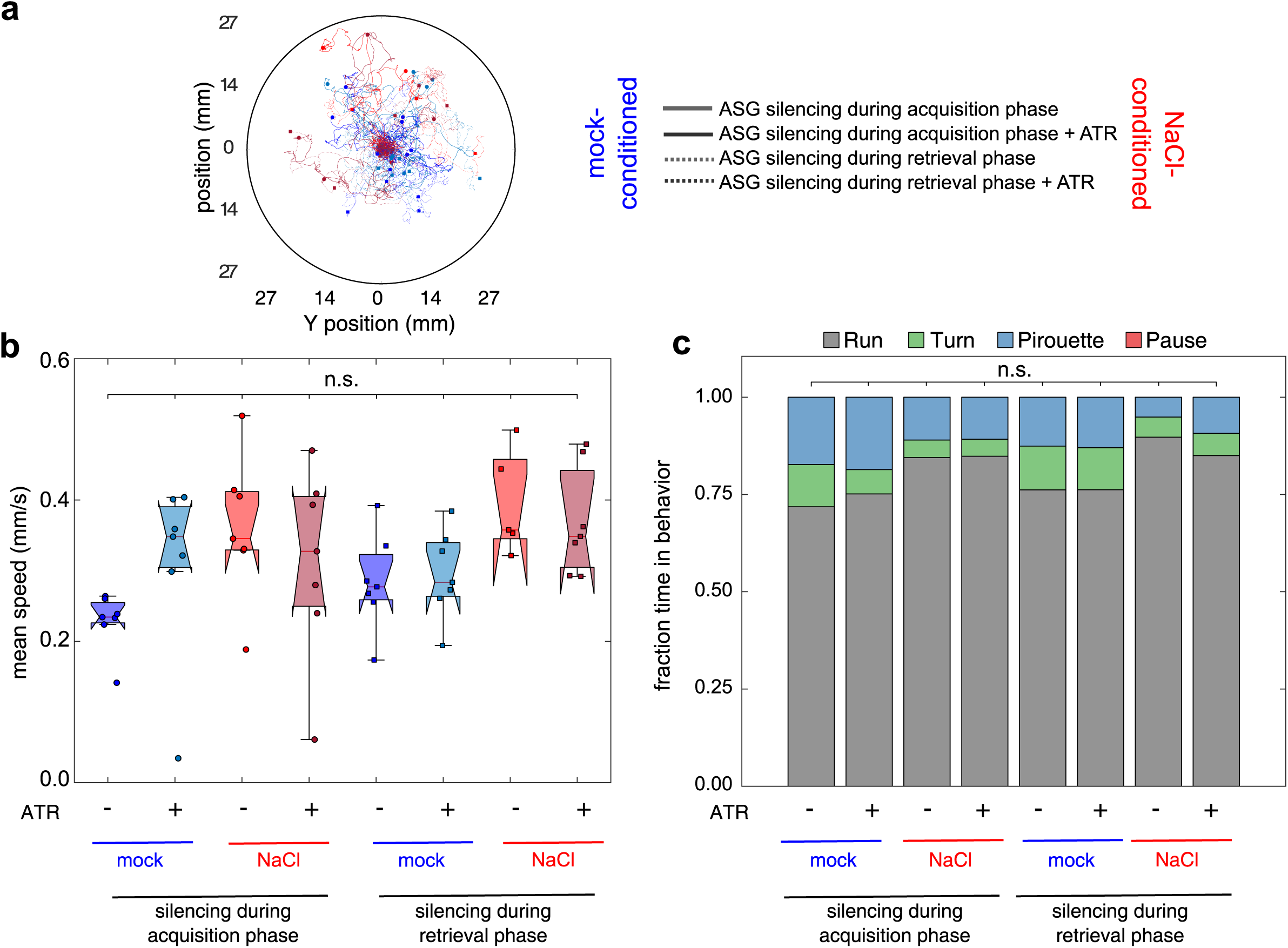
Behavioral effects of silencing CAPA-1 neurons (related to Figure 6) **(a)** The inhibitory opsin Arch is expressed in ASG under control of the *capa-1* promoter. Individual transgenic worms are either illuminated with yellow-green light for ASG silencing while being conditioned, or when they are navigating an agar plate without NaCl supplemented to it. Arch requires the cofactor *all-trans* retinal (ATR) to be supplemented to the worm culture. Transgenic *C. elegans* not fed ATR-supplemented food serve as a negative control. Representation of individual tracks on 55 mm diameter plates without NaCl added to it. Worms are released in the middle of the plate, after which they are kept in the field-of-view of the illumination microscope using an automated xy-stage. n ≥ 5 animals per genotype. **(b)** Mean speed of individual worms. **(c)** The fraction of time in each behavioral state is plotted for each genotype. Statistical comparisons by Chi-square test on binned data. n.s., not significant.

## Supplementary resource table

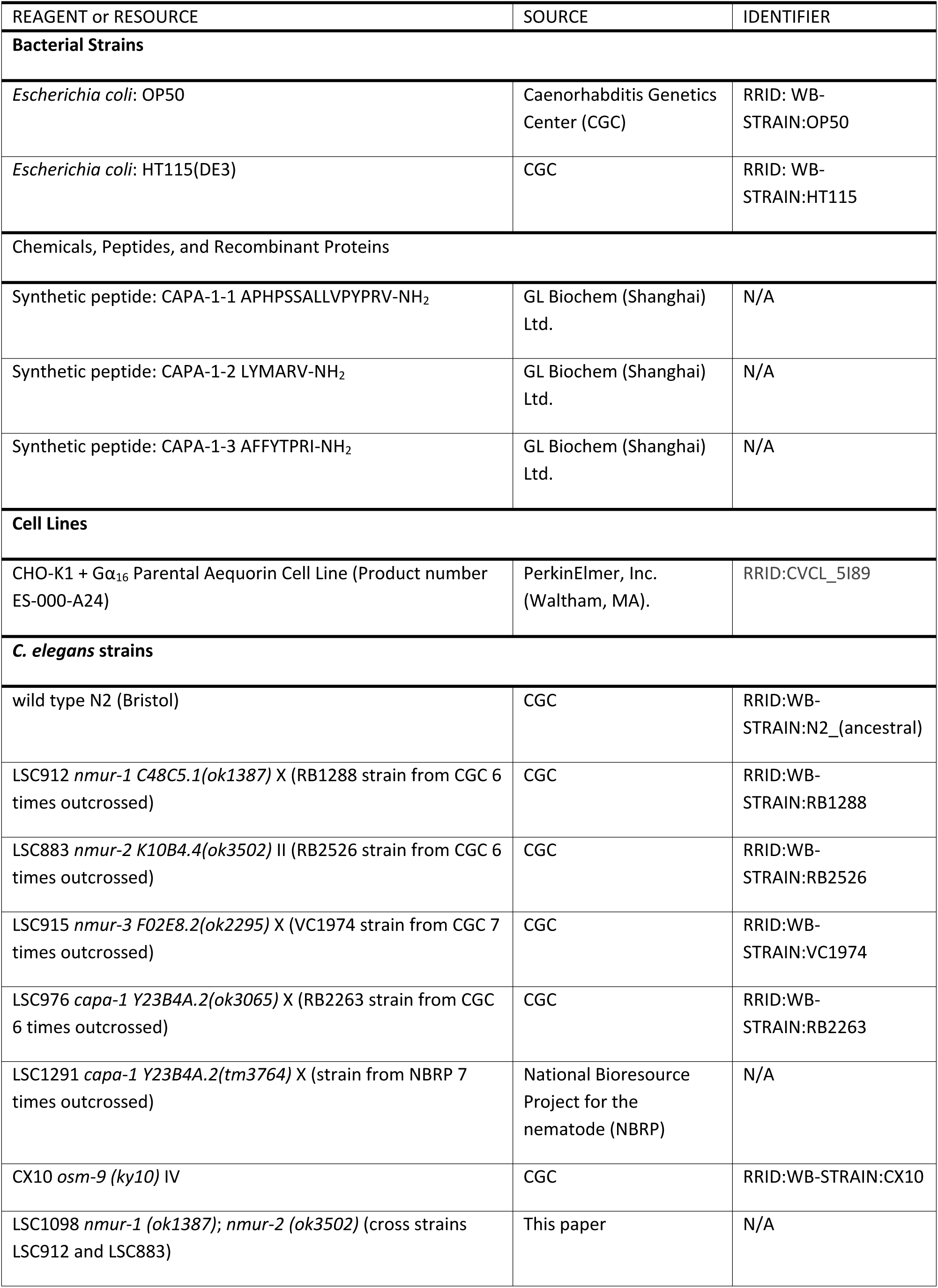
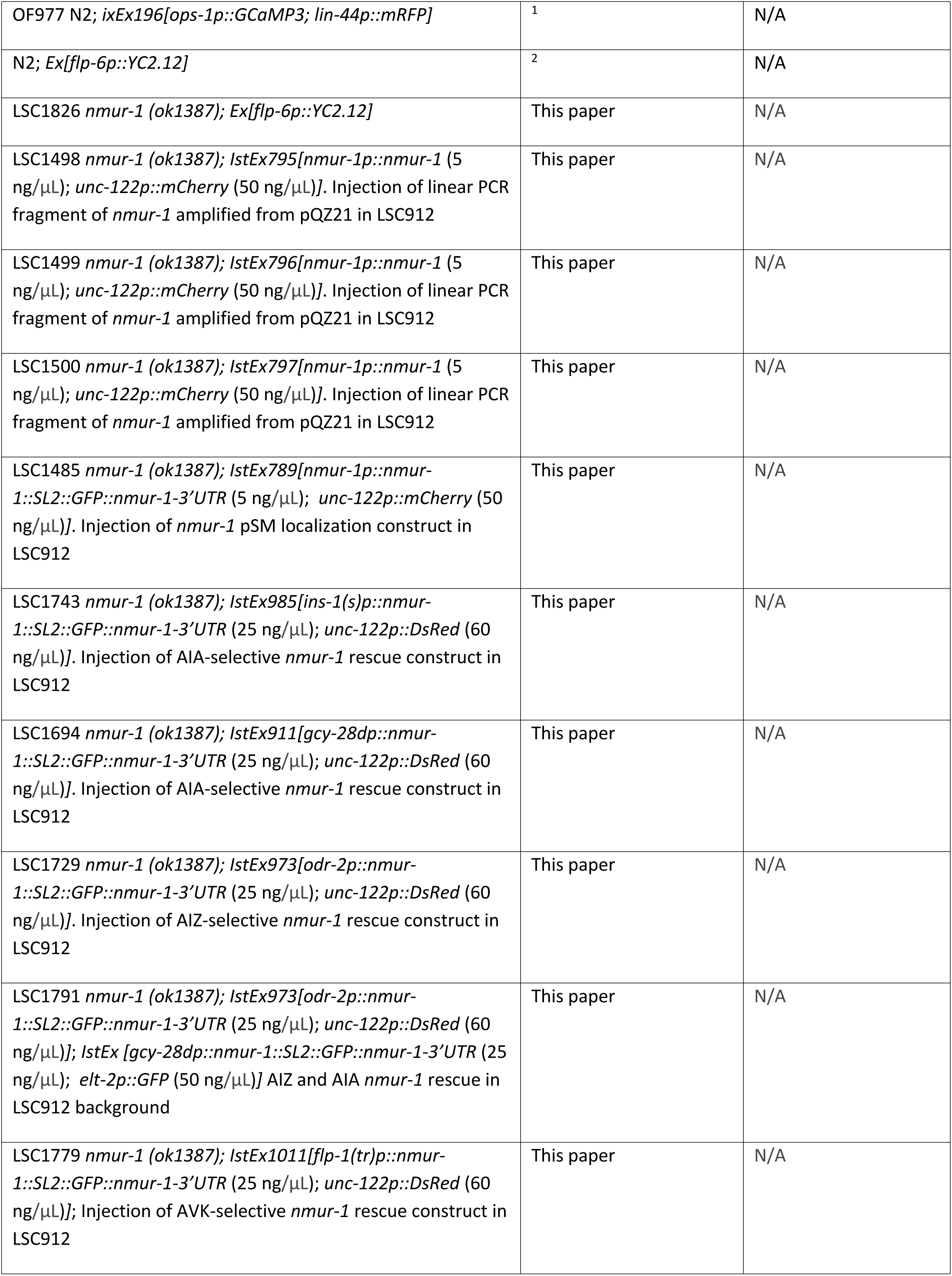
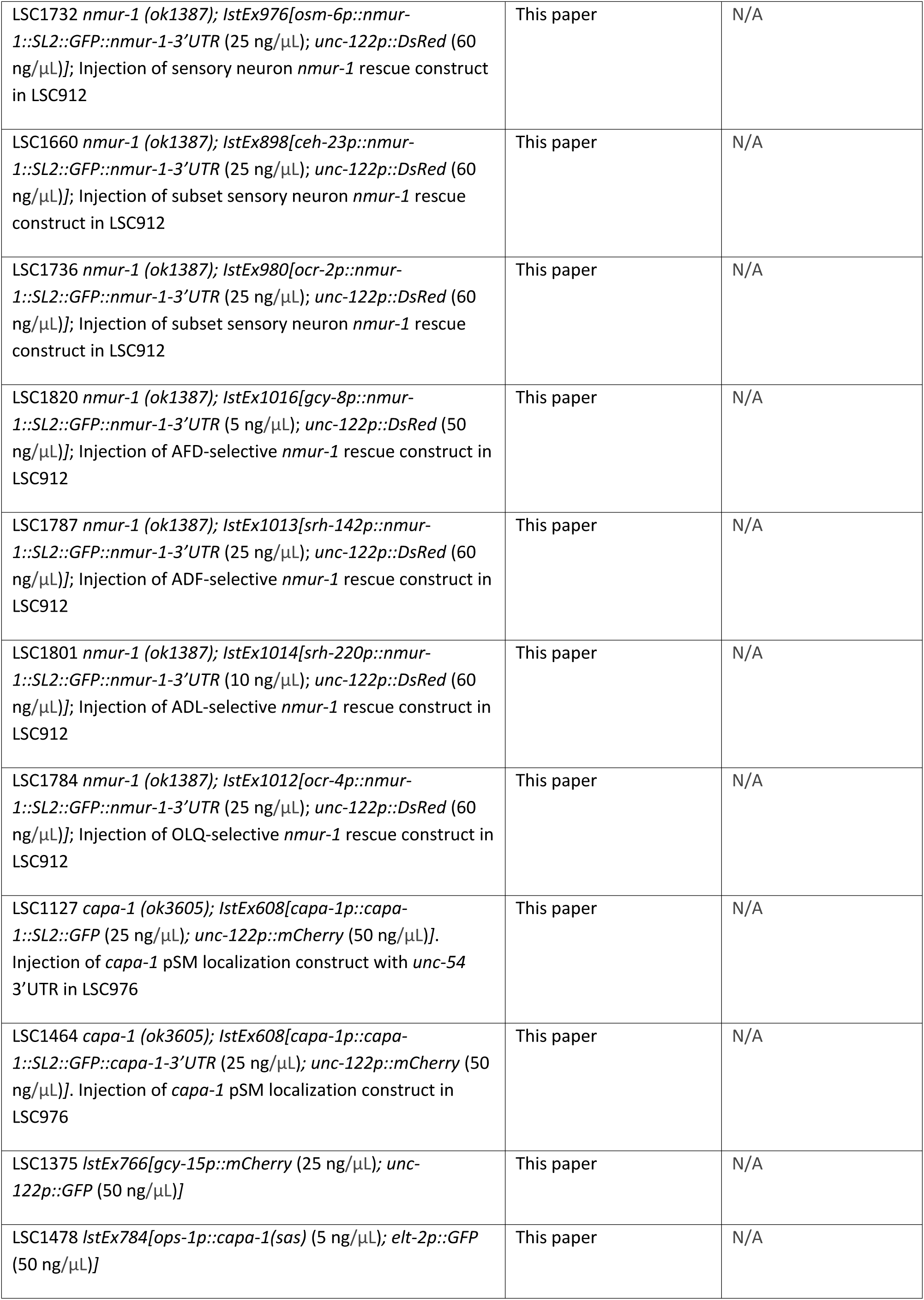
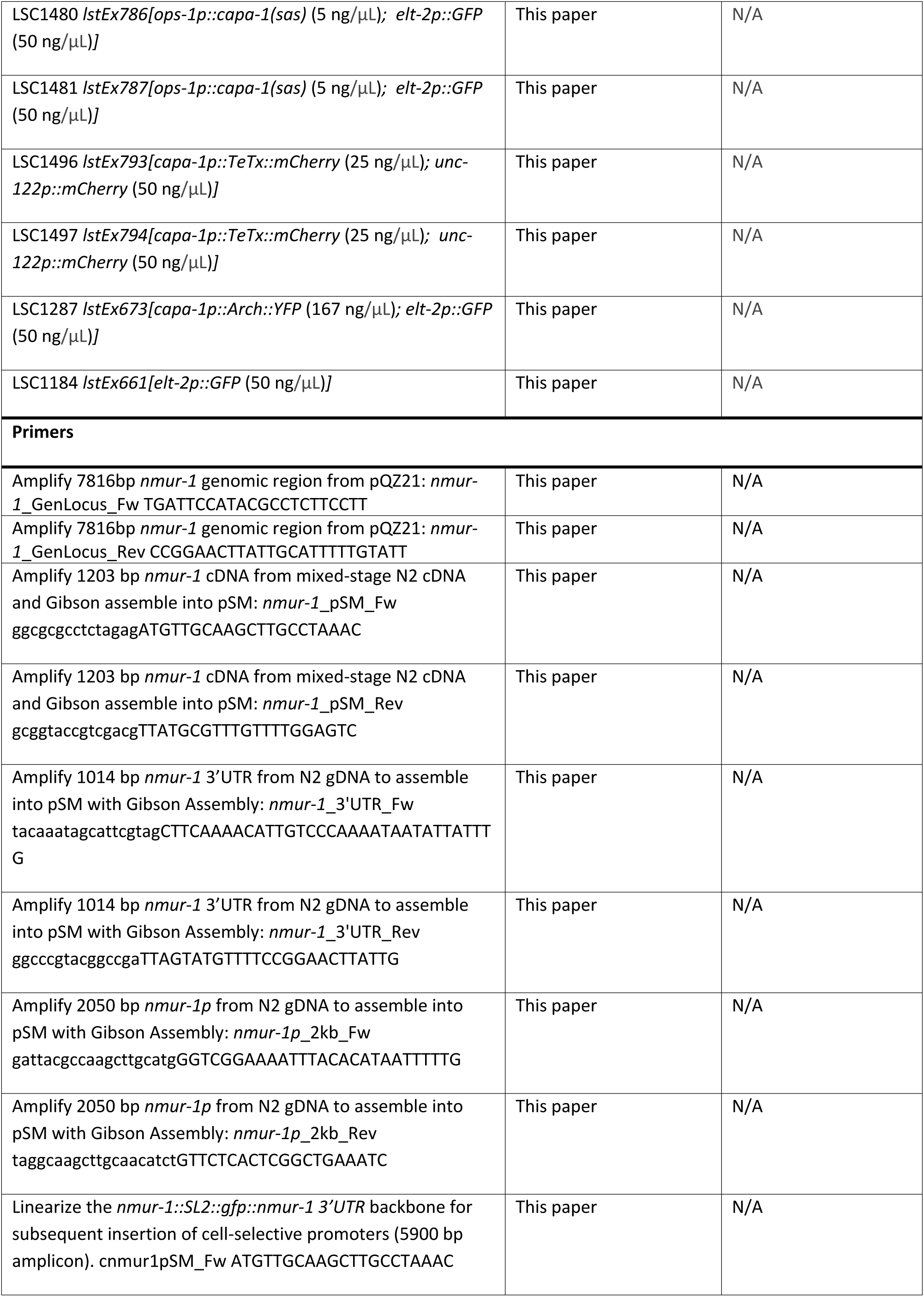
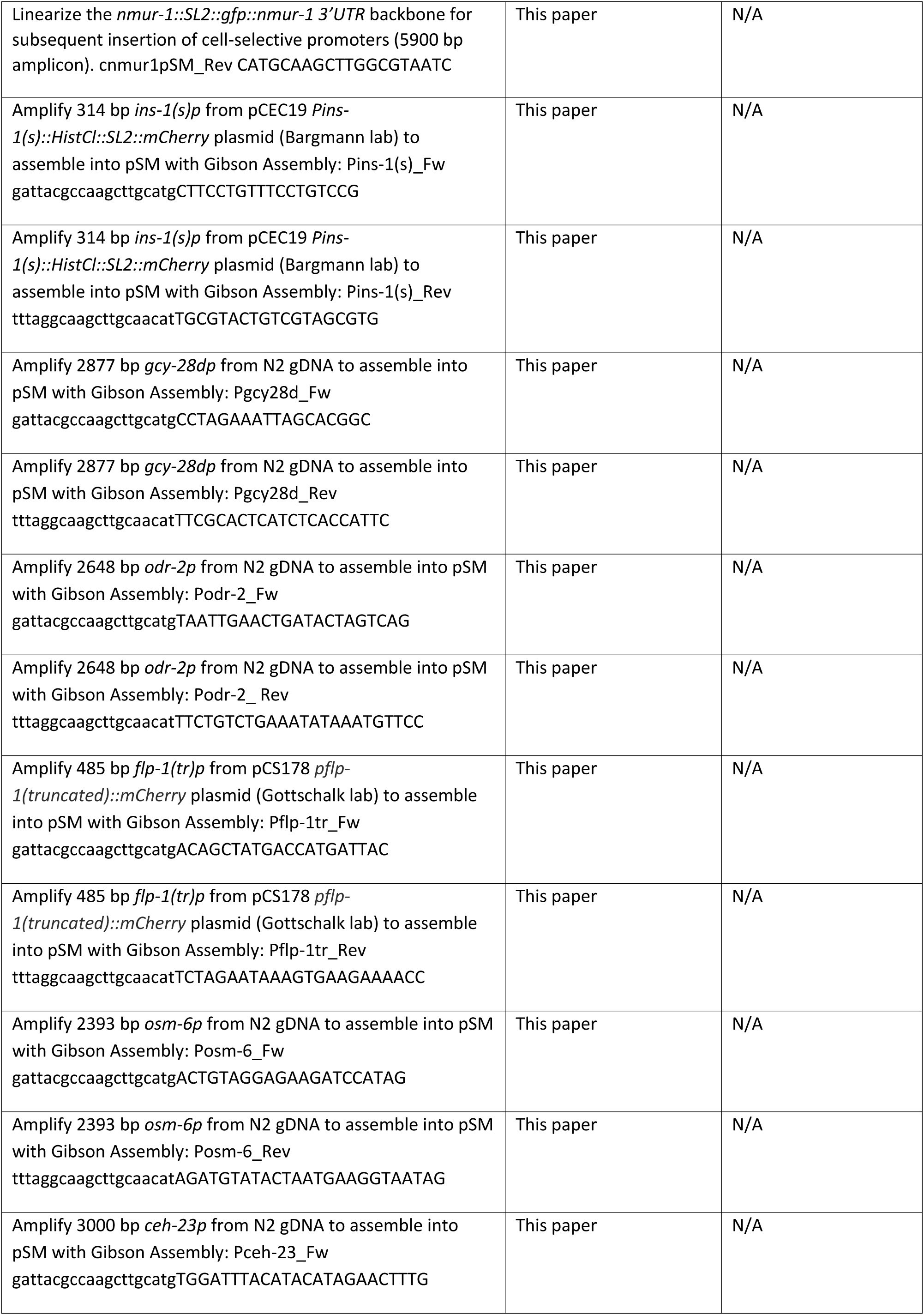
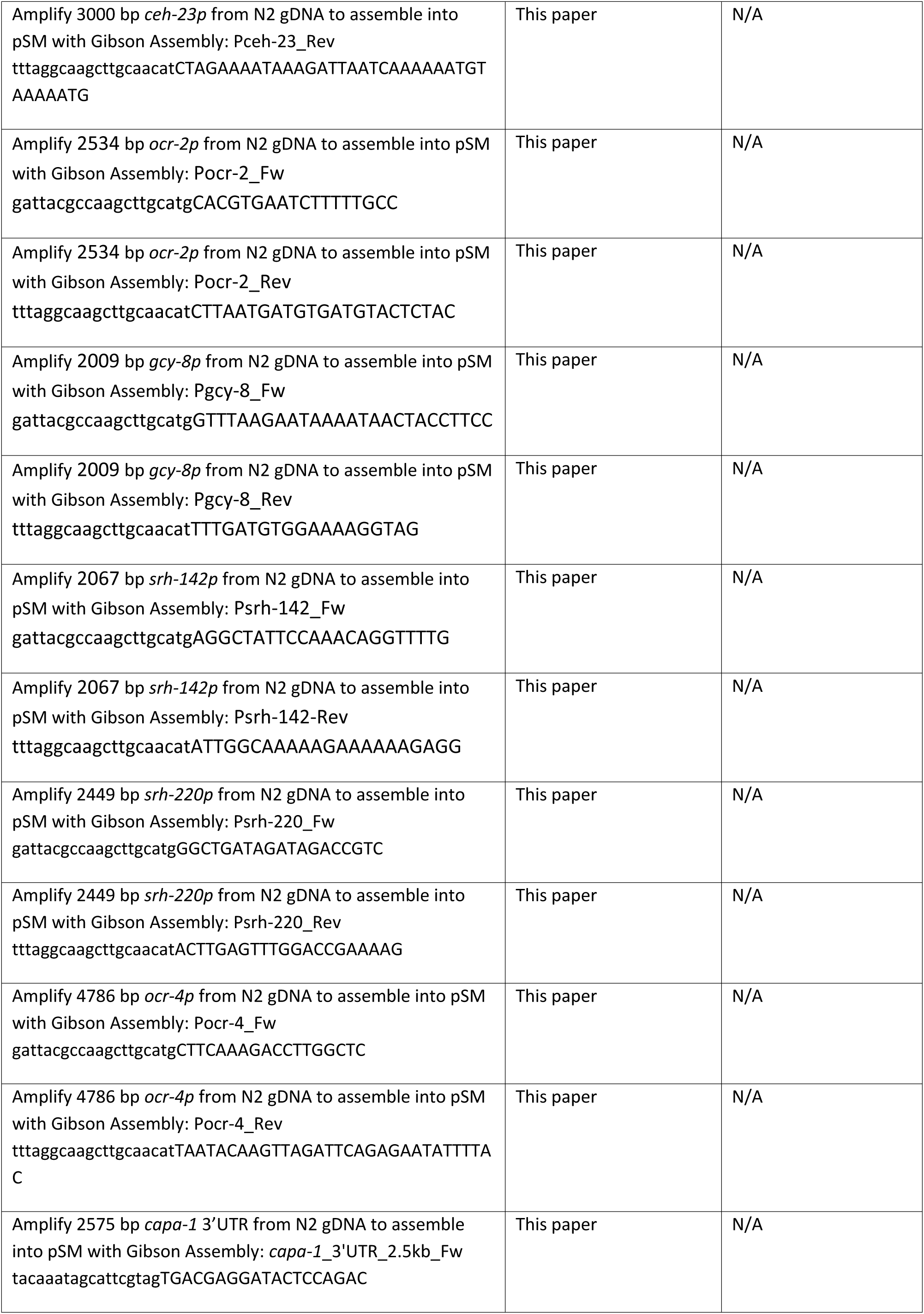
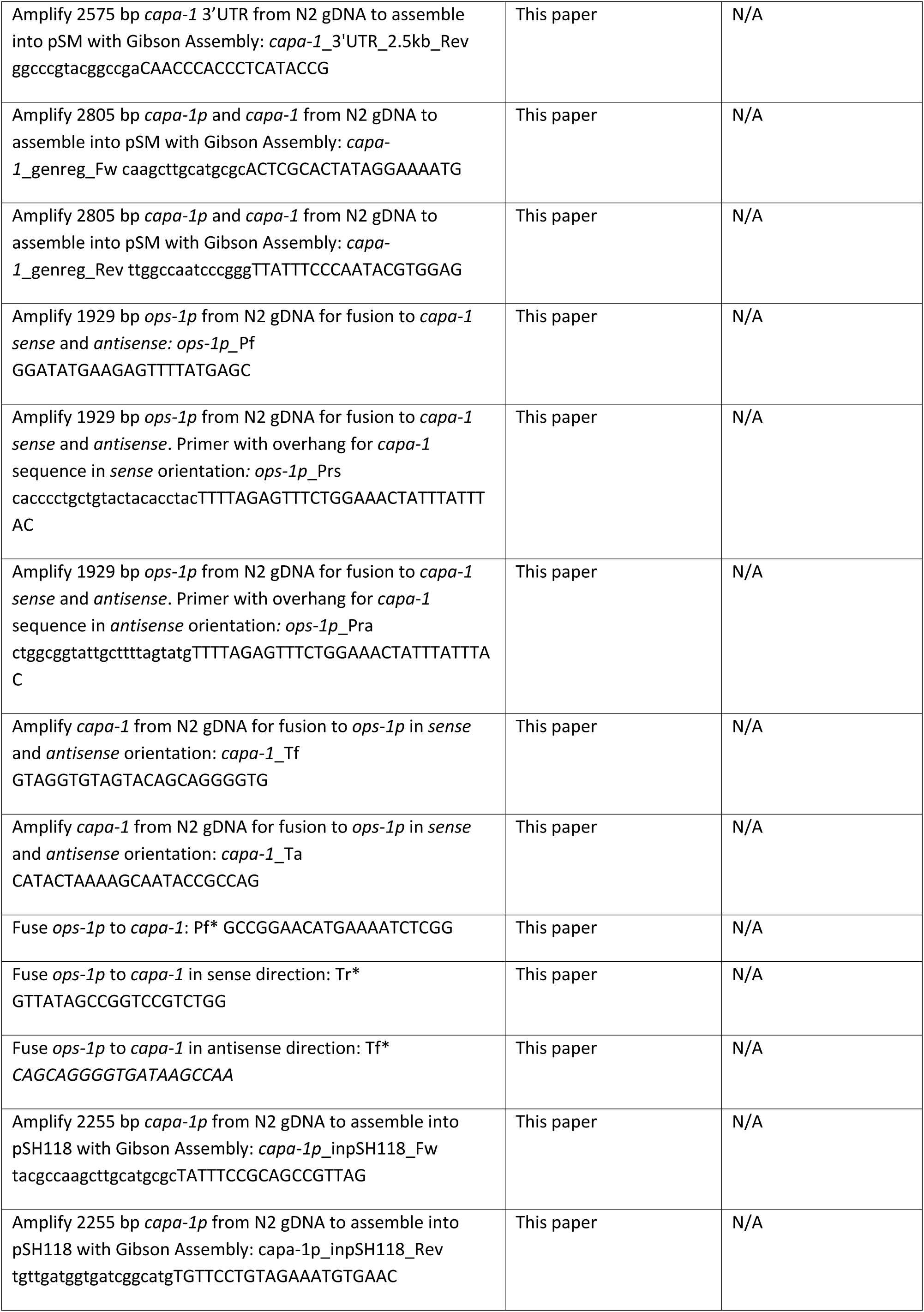
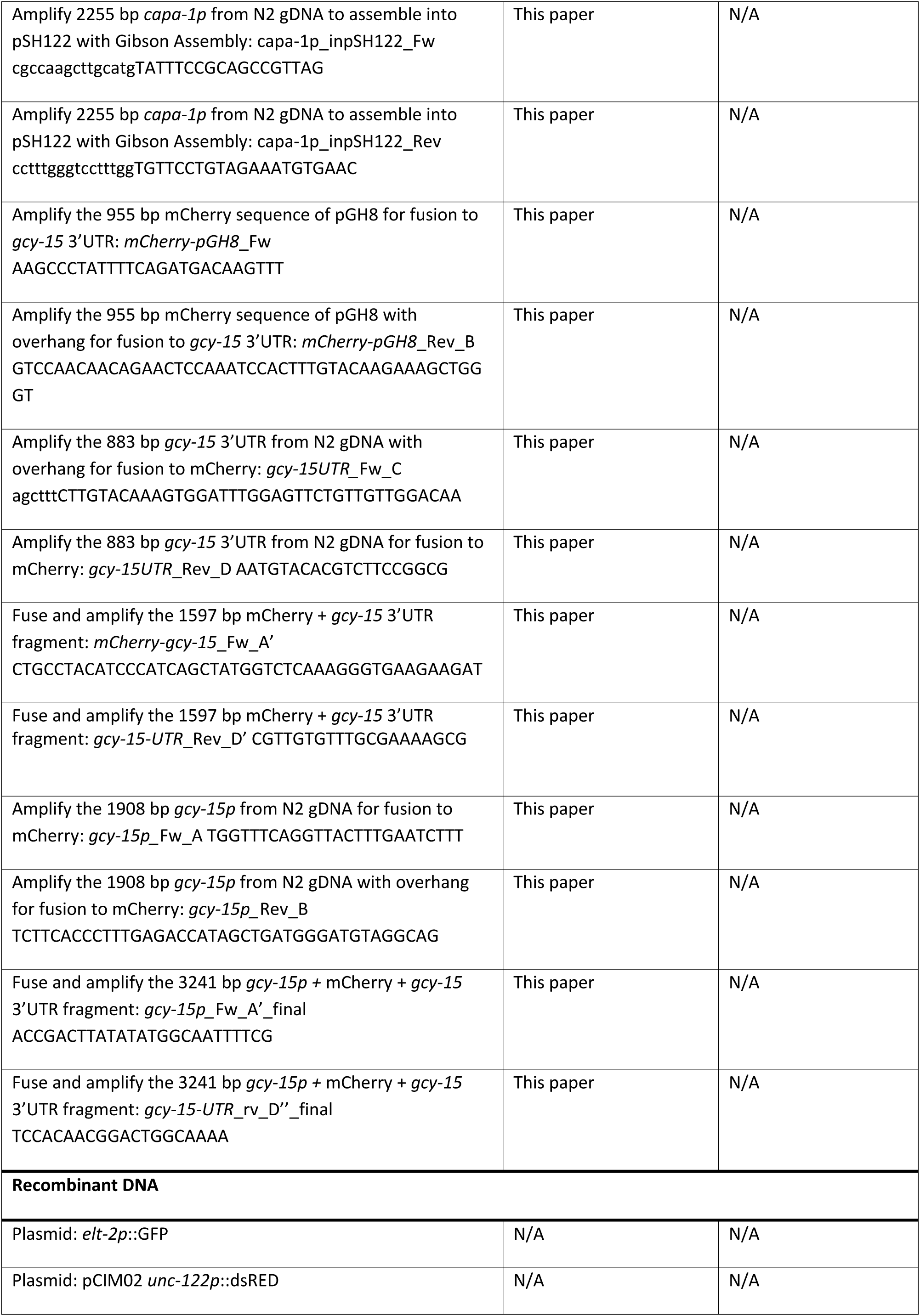
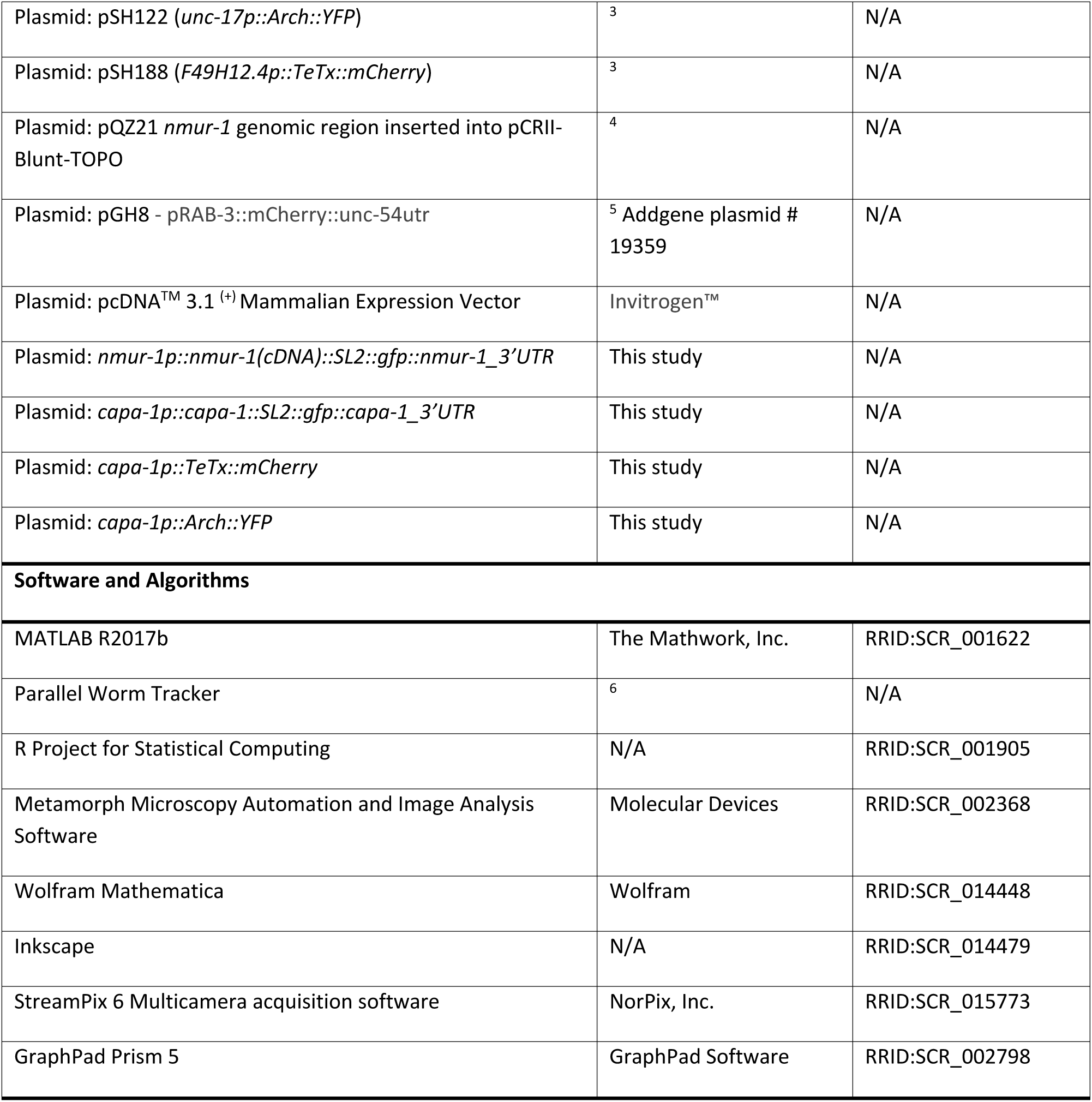

